# Organization and engagement of a prefrontal-olfactory network during olfactory selective attention

**DOI:** 10.1101/2021.09.04.458996

**Authors:** Hillary L. Cansler, Estelle E in ’t Zandt, Kaitlin S. Carlson, Waseh T. Khan, Minghong Ma, Daniel W. Wesson

## Abstract

Sensory perception is profoundly shaped by attention. Attending to an odor strongly regulates if and how a smell is perceived – yet the brain systems involved in this process are unknown. Here we report integration of the medial prefrontal cortex (mPFC), a collection of brain regions integral to attention, with the olfactory system in the context of selective attention to odors. First, we used tracing methods to establish the tubular striatum (TuS, also known as the olfactory tubercle) as the primary olfactory region to receive direct mPFC input in rats. Next, we recorded local field potentials from the olfactory bulb (OB), mPFC, and TuS while rats completed an olfactory selective attention task. Gamma power and coupling of gamma oscillations with theta phase were consistently high as rats flexibly switched their attention to odors. Beta and theta synchrony between mPFC and olfactory regions were elevated as rats switched their attention to odors. Finally, we found that sniffing was consistent despite shifting attentional demands, suggesting that the mPFC-OB theta coherence is independent of changes in active sampling. Together, these findings begin to define an olfactory attention network wherein mPFC activity, as well as that within olfactory regions, are coordinated in manners based upon attentional states.

## Introduction

Sensory processing and thus perception are both profoundly shaped by our ever-changing cognitive states. In most cases, the thalamus appears to be a major driver of state-dependent modulation of sensory information (Wimmer *et al*. 2015; O’Connor *et al*. 2002; Halassa and Kastner 2017; McCormick and Feeser 1990). For instance, the visual thalamus modulates primary visual cortex in manners that enhances the signal to noise of an attended visual cue (McAlonan *et al*. 2008). Similarly, the gustatory thalamus regulates taste-evoked responsivity of gustatory cortex neurons (Samuelsen *et al*. 2013). Among all sensory systems, the olfactory system presents a unique challenge for understanding the influence of cognitive state upon sensory processing. This is because, while humans (Zelano *et al*. 2005; Spence *et al*. 2000; Plailly *et al*. 2008) and rodents (Carlson *et al*. 2018) alike can selectively attend to odors, the organization of the olfactory system lacks obligatory thalamic processing (Gottfried 2010; Courtiol and Wilson 2016; Kay and Sherman 2006). Thus, other brain systems must engage with olfactory processing in order to afford one the ability to attend to odors. This is a significant issue since odors are most often encountered in highly multisensory environments, for instance during eating, wherein potentially distracting or conflicting cues must be ignored at the expense of selectively attending to odor.

Truly very little is known regarding the neural mechanisms underlying olfactory attention. One brain region that seems likely to confer this ability, at least in part, is the tubular striatum (TuS, also known as the olfactory tubercle (Wesson 2020)). This is true in both humans and rodents. For instance, early work using fMRI uncovered the first evidence that the human TuS is more activated in response to attended versus unattended odors while human subjects engaged in an olfactory selective attention task (Zelano *et al*. 2005). Importantly, attention-dependent amplification of odor-evoked activity in the TuS exceeded that of even the primary “piriform” olfactory cortex (PCX) (Zelano *et al*. 2005). Our group subsequently found that odor-evoked signal to noise among TuS neurons was enhanced as rats engaged in olfactory selective attention. (Carlson *et al*. 2018). While the TuS is engaged by attention in manners which may subserve the ability to attend to odors, the way that the TuS integrates into a wider brain network in the context of odor-directed attention is unknown. This includes major voids in our understanding of descending inputs from brain regions known to be integral for attention, and how TuS activity is structured relative to that of these regions.

The rodent prefrontal cortex (PFC), depending upon how one chooses to define it (Laubach *et al*. 2018; Le Merre *et al*. 2021), comprises several key subregions including but not limited to the medial PFC (mPFC) and the orbitofrontal cortex (OFC), each of which can be divided into more specific subregions. The medial prefrontal cortex (mPFC) is crucial for many executive processes including attention (Wimmer *et al*. 2015; Birrell and Brown 2000; Kim *et al*. 2016; Miller and Cohen 2001) and is highly interconnected with the rest of the brain (Le Merre *et al*. 2021), with particularly dense inputs to sensory and thalamic areas. mPFC neurons are modulated as animals attentively await a stimulus (Rodgers and DeWeese 2014; Kim *et al*. 2016), and disruptions to the mPFC impair attentional set-shifting (Birrell and Brown 2000; Ragozzino *et al*. 2003), sustained attention (Kim *et al*. 2016), and selective attention (Wimmer *et al*. 2015). mPFC subdivisions include the prelimbic (PrL) and infralimbic (IL) cortices, which seem to possess dissociable behavioral functions (Hardung *et al*. 2017; Marquis *et al*. 2007; de Kloet *et al*. 2021; Luchicchi *et al*. 2016). Specifically, the PrL appears important for set-shifting and selective attention (Marquis *et al*. 2007; Schmitt *et al*. 2017; Kim *et al*. 2016; Rodgers and DeWeese 2014), while the IL is implicated in behavioral flexibility and extinction (Barker *et al*. 2014). Therefore, the mPFC, especially the PrL and IL, are putative candidates for influencing olfactory processing via top-down modulation during attentional states.

Local field potential (LFP) oscillations in the gamma band (40-100 Hz) in sensory and prefrontal cortices have been associated with attention (Fries *et al*. 2001; Vinck *et al*. 2013; Borgers *et al*. 2005; Brassai *et al*. 2015; Schroeder and Lakatos 2009a). In the olfactory system, increased power of gamma oscillations in the olfactory bulb (OB) is observed during successful discriminations between perceptually demanding odor pairs (Beshel *et al*. 2007), and learning (Losacco *et al*. 2020a), suggesting that they in some manner aid in cognitively demanding processes. In addition to high frequency oscillations, theta oscillations (2-12Hz) are profoundly shaped by respiration, including fast investigatory sniffing, in olfactory regions and beyond (Adrian 1942; Macrides *et al*. 1982; Vanderwolf 1992; Tort *et al*. 2018b; Colgin 2013; Zhang *et al*. 2021; Fontanini and Bower 2006; Kay and Laurent 1999; Buonviso *et al*. 2003; Miura *et al*. 2012). Interestingly, theta-band coherence is elevated between the OB and mPFC in emotionally-salient contexts (Moberly *et al*. 2018; Bagur *et al*. 2021; Zhong *et al*. 2017), indicating functional connectivity between these networks that can be influenced by behavioral state. Examining neural oscillations within individual structures, and their synchrony between structures, can yield valuable insights into the ways that brain regions form functional networks (Buzsaki 2006; Fries 2015). Whether the mPFC and olfactory system networks (either independently or together) are engaged by selective attention to odors has not been explored.

Here, we sought to investigate the anatomical and functional integration of the mPFC with olfactory regions (including the OB and TuS) in the context of selective attention to odors. First, we used cell-type-specific tracing methods to reveal that excitatory mPFC neurons preferentially target the TuS compared with other olfactory regions. Next, we used multi-site LFP recordings to demonstrate that the OB, mPFC, and TuS modify their activity in intra- and inter-areal manners during a behavioral task that requires selective attention to odors and intermodal attentional switches. Interestingly, through measuring sniffing as rats engaged in selective attention, we observed that rats covertly display odor-directed attention, maintaining highly stereotyped sniffing structured to the task despite shifting attentional demands. Together this work adds to our understanding of the organization and activities of brain systems which are engaged during olfactory attention.

## Results

### The PrL and IL preferentially target the TuS compared to other major olfactory structures

The rodent PFC projects throughout the brain with notably strong and well characterized inputs to the ventral striatum and thalamus (Vertes 2004; Le Merre *et al*. 2021). Yet the connectivity of the PFC with primary (OB) and secondary olfactory structures (the anterior olfactory nucleus (AON), PCX, and TuS) is not well defined. To address this, we injected Cre-dependent anterogradely transported AAVs encoding synaptophysin tagged with either GFP or mRuby into the PrL or IL, in combination with an AAV encoding Cre under control of the CaMKII promotor **(Fig. 1A-B)** (Herman *et al*. 2016). This approach allowed us to observe fluorescent puncta (which can more confidently be attributed to synaptic terminals, rather than fibers of passage) in olfactory regions receiving input from either the PrL or IL cortex. Importantly, because Cre expression was driven by the CaMKII promotor, we can specifically assess excitatory projections which make up >80% of mPFC neurons (Erö *et al*. 2018) and are major regulators of the PFC’s effects (de Kloet *et al*. 2021). We quantified puncta in three structures recipient of dense olfactory bulb input: the AON, PCX and TuS **(Fig. 1B-G)**. We also inspected the OB for puncta, but did not quantify this since none were detectable **(Fig. 1F)**. Other than the OB, we observed fluorescent puncta in all regions examined, with a striking density in the TuS. Specifically, the medial division of the TuS (mTuS), which some have indicated plays a particularly prominent role in olfaction and motivated behaviors (Ikemoto 2003; Murata *et al*. 2015; Zhang *et al*. 2017), receives the most input from the mPFC, even more so than all other regions combined **(Fig. 1G)**. Within the mTuS, we observed synapses throughout all 3 layers, with significantly more in layers 2 and 3 **(Fig. 1H)**. Importantly, this is where the vast majority of medium spiny neurons, the principal neuron of the TuS, reside. Further, we found that the IL projections to the mTuS are denser than those of the PrL **(Fig. 1G-H)**. Notably, in 2 separate rats we unilaterally injected a retrograde AAV encoding GFP into the PrL and IL and observed no GFP+ cells in the TuS (or anterior PCX), indicating that the there is no direct reciprocal feedback from the TuS (or anterior PCX) to the PrL or IL (**Fig. S1**). Together, these findings indicate that the PrL and IL project to multiple olfactory regions, with the mTuS being the primary recipient of this input.

**Figure 1.**
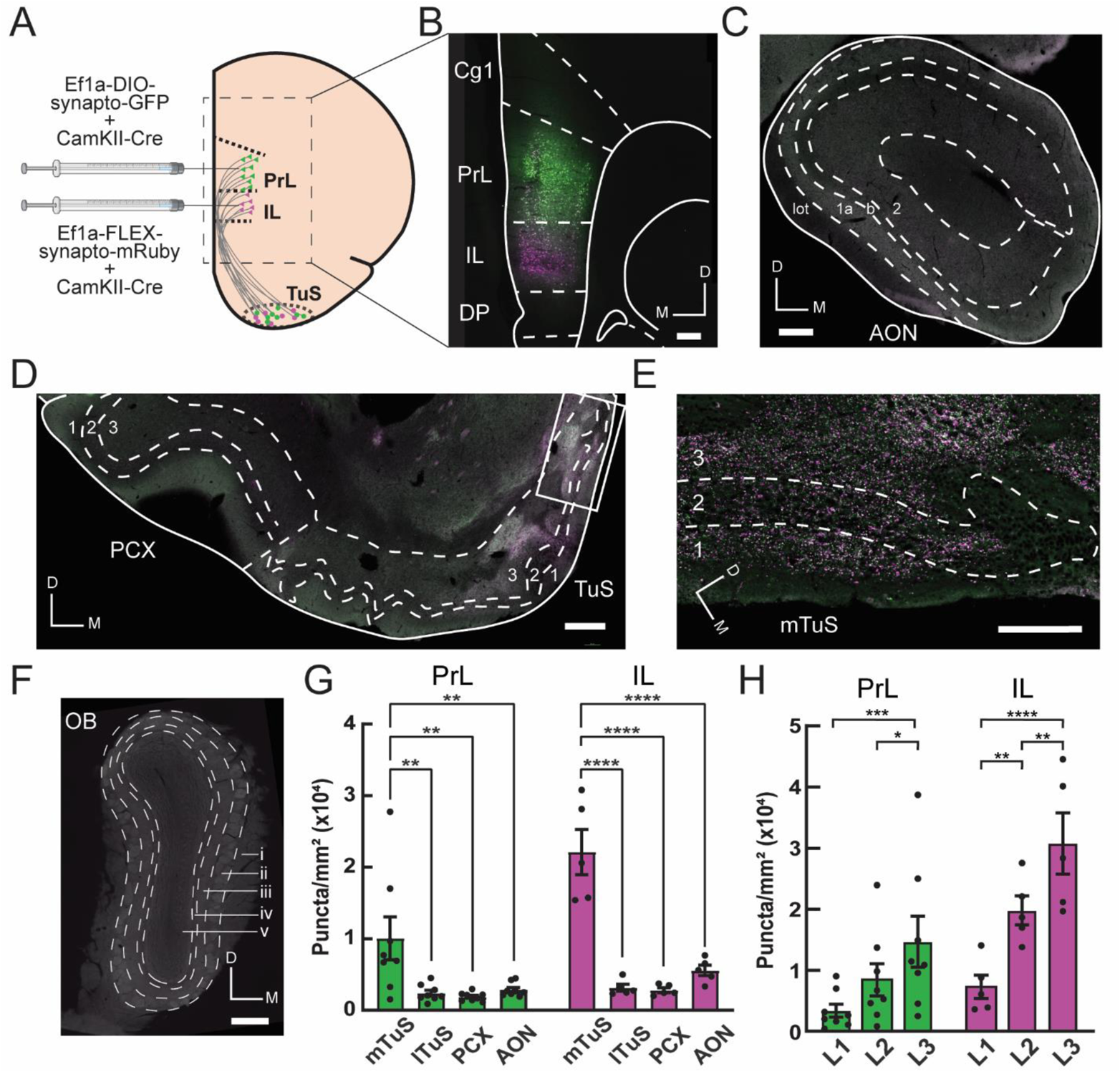
The prelimbic and infralimbic medial prefrontal cortex preferentially target the tubular striatum compared to other major olfactory structures. **A**. The PrL and IL cortices were selectively targeted with 50/50 mixtures of Ef1a-DIO-synaptophysin-GFP/pENN-AAV9-CamKII-Cre-SV40 and Ef1a-FLEX-synaptophysin-mRuby/pENN-AAV9-CamKII-Cre-SV40, respectively. **B**. Representative mPFC image showing region-specific viral transduction within the same rat. Scale bar 250 μm. **C**. Representative image of the AON, showing few fluorescent puncta. Cell layers 1-2 and the lateral olfactory tract (lot) are indicated. Scale bar 250 μm. **D**. Representative image of the PCX and TuS. Note high fluorescence in the medial TuS and low fluorescence in the lateral TuS and PCX. Boxed region is shown in panel E. Scale bar 250 μm. **E**. Magnified view of boxed region shown in panel D, showing high levels of fluorescent puncta, indicating synaptic terminals from TuS-projecting mPFC neurons. This image has been digitally deconvolved to enhance clarity, for illustration purposes only. Scale bar 100 μm. **F**. Representative image of the OB absent of fluorescent puncta. Dashed lines indicate layers: i. olfactory nerve layer; ii. glomerular layer; iii. external plexiform layer; iv. mitral cell layer; v. granule cell layer. Scale bar 250 μm. **G**. Quantification of fluorescent puncta across olfactory regions, normalized by area of quantified region. PrL:one-way ANOVA, main effect of regions, F(3,21)=7.82, p=0.001. IL:one-way ANOVA, main effect of regions, F(3,12)=37.08, p<0.0001. Asterisks indicate results from Tukey’s multiple comparisons test, **p<0.01, ****p<0.0001. PrL mTuS vs. IL mTuS, unpaired t-test, p=0.02. H. Quantification of fluorescent puncta across layers in the mTuS. PrL: one-way ANOVA, main effect of layers, F(2,14)=12.62, p=0.0007. IL: one-way ANOVA, main effect of layers, F(2,8)=43.04, p<0.0001. Asterisks indicate results from Tukey’s multiple comparisons test, *p<0.05, **p<0.01, ***p<0.001, ****p<0.0001. PrL, prelimbic cortex; IL, infralimbic cortex; Cg1, cingulate area 1; DP, dorsal peduncular cortex; mTuS/lTuS, medial/lateral tubular striatum; PCX, piriform cortex; AON, anterior olfactory nucleus; OB, olfactory bulb; D, dorsal; M medial; L1-L3, layer 1-3. PrL injection, n=8 rats; IL injection, n=5 rats. All error bars represent SEM.

### Among PFC subregions, layer 5 PrL and IL neurons provide the densest input to the TuS

While the PrL and IL are strongly implicated in attention, the PFC also includes the OFC. The OFC is involved in polysensory processing (Rolls 2004; de Araujo *et al*. 2003; Small *et al*. 2001), and cognition and decision-making (for review see (Izquierdo 2017; Schoenbaum *et al*. 2009)), making it another strong candidate circuit to instruct state-dependent odor processing. To directly compare the OFC→TuS and mPFC→TuS pathways, we unilaterally injected a retrograde AAV encoding GFP into the TuS, and quantified cell bodies throughout the PFC, including the PrL, IL, medial (MO), ventral (VO), lateral (LO) and dorsolateral (DLO) OFC subdivisions **(Fig. 2A-D)**. We found the greatest numbers of cells in the IL, PrL, and MO, which together make up the ventromedial PFC (vmPFC) by virtue of their similar connectivity patterns (Le Merre *et al*. 2021). We observed significantly more cells in the IL than all other regions quantified except the PrL, and significantly more cells in the PrL than all other areas regions quantified except the IL and the MO **(Fig. 2D)**. Thus, the PrL and IL provide the densest inputs to the TuS and are well-positioned to influence odor processing.

**Figure 2.**
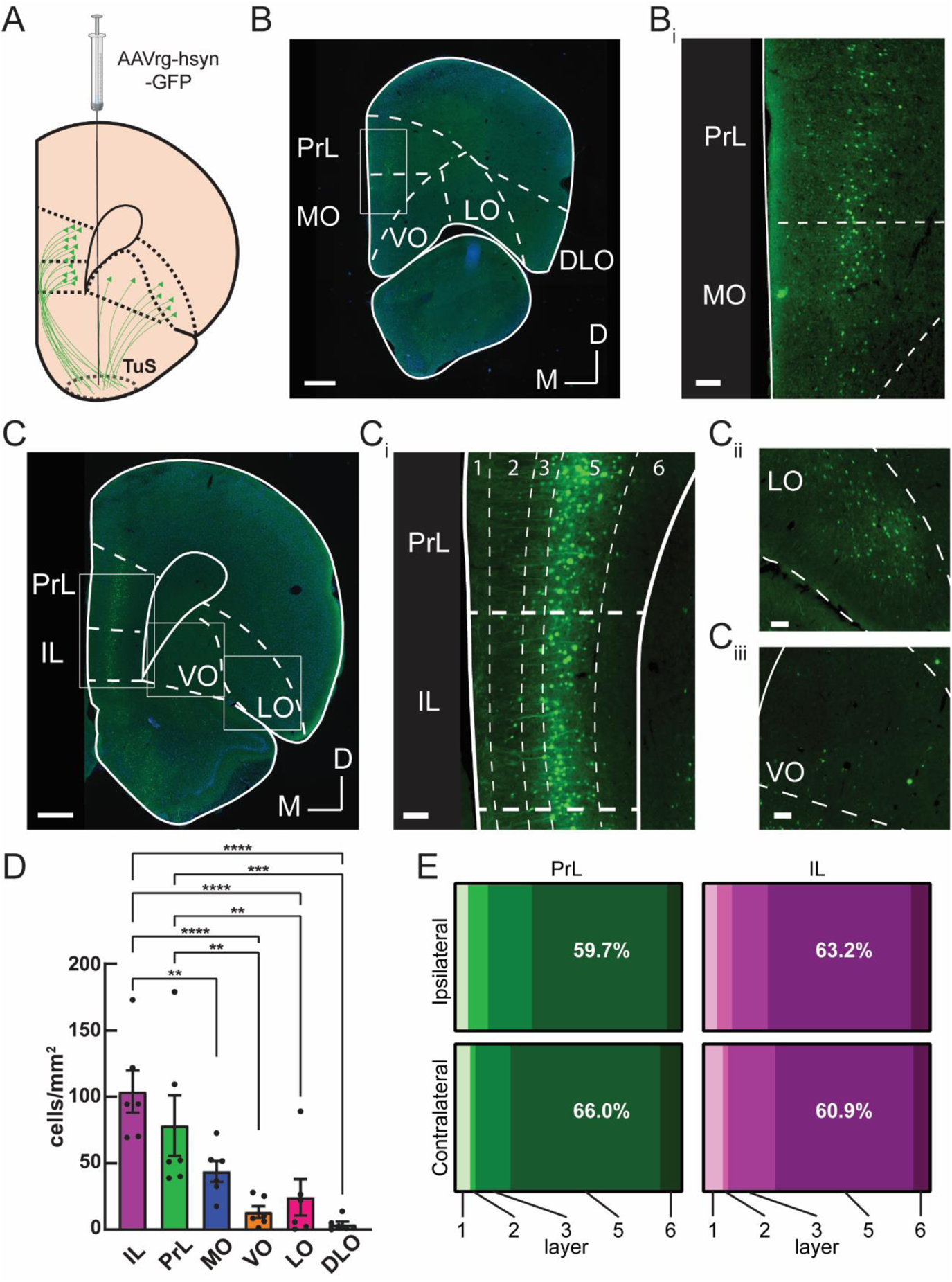
Among prefrontal cortex subregions, layer 5 prelimbic and infralimbic neurons provide the densest input to the tubular striatum. **A**. The TuS was injected with AAVrg-hsyn-GFP to identify TuS-projecting neurons throughout the prefrontal cortex. **B**. Representative mPFC image at Bregma +4.2mm, showing GFP-labeled TuS-projecting neurons. Boxed region is indicated in panel B_i_. Scale bar 500 μm. **B_i_**. Magnified view of the boxed region in panel B_i_. Scale bar 100 μm. **C**. Representative PFC image at Bregma +3.2 mm, showing GFP-labeled TuS-projecting neurons. Scale bar 500 μm. **Ci**. Magnified view of boxed region in panel C showing the PrL and IL cortices. Dotted lines indicate layers. Scale bar 100 μm. **Cii**. Magnified view boxed region showing LO cortex. Scale bar 100 μm. **Ciii**. Magnified view of boxed region showing VO cortex. Scale bar 100 μm. n=6 rats. **D**. Quantification of cell numbers across prefrontal cortex regions ipsilateral to the injection site. One-way ANOVA, main effect of regions, F(5, 25)=15.67, p<0.0001. Asterisks indicate results of Tukey’s multiple comparisons test, **p<0.01, ***p<0.001, ****p<0.0001. Error bars represent SEM. **E**. Distribution of cell bodies across PrL and IL layers, in both the contralateral and ipsilateral hemispheres, showing the majority of cell bodies are found in layer 5. PrL: Two-way ANOVA, main effect of layer F(1.15, 5.76)=11.48, p=0.014. IL: Two-way ANOVA, main effect of layer F(1.32, 6.61)=42.55, p=0.0003; main effect of hemisphere F(1,5)=34.39, p=0.002. n=6 rats.

Within the PrL and IL, we quantified cell body locations throughout the cortical layers for both the ipsilateral and contralateral hemispheres. We observed that in both of these regions, the majority of cell bodies were found in layer 5 **(Fig. 2E)**, which is consistent with other glutamatergic (de Kloet *et al*. 2021) corticostriatal mPFC projections (Nakayama *et al*. 2018; Ding *et al*. 2001; Gabbott *et al*. 2005). The fact that the TuS receives its densest PFC inputs from the PrL and IL cortices, together with the fact that these regions preferentially target the TuS over other olfactory regions, indicates that the mPFC→TuS pathway is likely the primary route whereby the olfactory system might receive information regarding states or tasks requiring high executive function, including attention.

### Investigating mPFC and olfactory network activity during odor-directed selective attention

We next sought to functionally test whether the mPFC and the olfactory system, specifically the OB and the TuS, integrate into a network during olfactory attention. To accomplish this, we combined multisite LFP recordings along with the Carlson Attention Task (CAT) (Carlson *et al*. 2018) to manipulate selective attention to odors **(Fig. 3)**. Briefly, the CAT is a modified two-alternative choice task in which rats are simultaneously presented with one of two olfactory cues (odor A/odor B) and one of two auditory cues (tone on/tone off). A single behavioral session begins with tone attention: that is, the tone cues signal the reward port location whereas the odor cues are distractors (**Fig. 3A-C**, blue shading). Once criterion on the tone attention phase of the task has been reached (6 blocks of 20 trials at ≥80% correct), an uncued intermodal rule change occurs, and the rats must now direct their attention to odors, and ignore tones, to accurately locate their rewards (**Fig. 3A-C**, orange shading). This rule change is accompanied by a temporary drop in performance as the rats adjust their behavior to the new rule (**Fig. 3C**, pink shading) before they eventually perform well on odor attention (6 blocks at ≥80% correct, orange shading). We will refer to the blocks following the rule change before performance reaches ≥80% correct on odor attention as “switch” blocks, in which the rat is by-definition performing poorly. Importantly, each session began and ended with 3 blocks of odor-only trials, in which there were no competing tone cues, to use as a control for odor discrimination without any additional cognitive demand.

**Figure 3.**
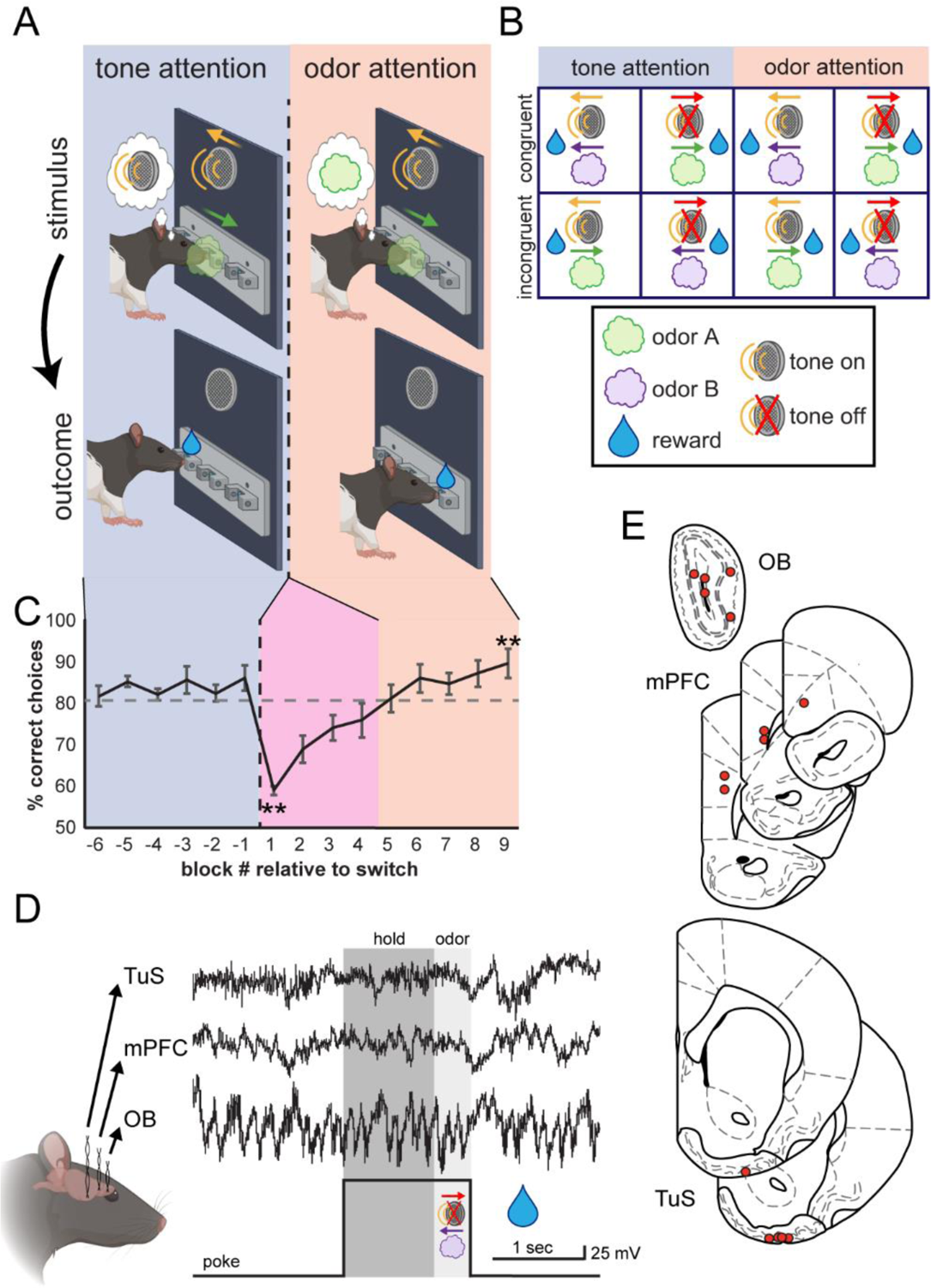
Investigating medial prefrontal cortex and olfactory network activity during odor-directed selective attention. **A**. Freely-moving rats initiate a trial by nose-poking in a center port, which triggers simultaneous delivery of one of two auditory cues and one of two odors (stimulus). These cues direct the rat to retrieve a fluid reward at either the left or the right port (outcome). Behavioral sessions begin with tone attention (auditory cues predict reward; blue shading) and switch to odor attention (odors predict reward; orange shading). **B**. All possible trial combinations in the Carlson Attention Task. Half of these are congruent (odor and tone indicate same reward port) and half are incongruent (odor and tone indicate opposite reward ports). **C**. Behavioral performance of all rats across behavioral sessions. After completing 6 blocks of tone attention at criterion (≥ 80% correct; blue shading), the task was switched to odor attention (orange shading). Rats then switched their attention to odors and completed 6 blocks at criterion. Block -1 vs. 1 paired, two-tailed t-test, **p = 0.001. Block 1 vs. 9 paired, two-tailed t-test, **p = 0.003. n=5 rats, 4.6 +/− 0.5 sessions per rat. Error bars represent SEM. **D**. All rats were implanted with bipolar recording electrodes in the OB, TuS, and mPFC, and LFPs were acquired during behavior. A sample trace is shown from a single trial, in which the rat pokes, holds in the center port for 1 second awaiting stimuli (dark gray shading), and remains for 400 ms to sample the stimuli (light gray shading). **E**. Electrode location summary. Red dots indicate tips of bipolar LFP electrodes.

In the CAT, there are four possible combinations of trials **(Fig. 3B)**. Two of these are “congruent,” in that the olfactory and auditory cues signal approach to the same reward port, and two are incongruent, in that the cues signal opposite ports. Thus, the correct reward port on incongruent trials depends on the current task rule: tone attention (blue shading) or odor attention (orange shading). Importantly, all analyses of physiological signals were limited to trials on which the tone was off to avoid multisensory influences and focus on the effects of cognitive state on odor processing specifically (Carlson *et al*. 2018). On a single trial of the CAT, the rat will nose poke to initiate a trial, then must hold in the center port for 1 second before the stimuli come on (**Fig. 3D**, dark gray shading). Then, the odor and tone stimuli come on simultaneously, and the rat must remain in the center port for at least 400 ms sampling the stimuli (**Fig. 3D**, light gray shading). After 400 ms, the rat is free to make a choice at the left or right port and receive a water reward if correct.

We simultaneously recorded LFPs from the TuS, mPFC, and OB from 5 highly-proficient expert rats (see Methods) while they performed the CAT **(Fig. 3D-E)**. This allowed us to explore network dynamics locally within each structure, as well as coherent activity between these structures.

### Elevations in gamma power upon intermodal switching and selective attention to odors

Gamma oscillations are widely observed throughout the brain, and are tied to a diverse array of functions including perception, memory, and attention (Fries *et al*. 2001; Buzsáki and Wang 2012; Mably and Colgin 2018; Kim *et al*. 2016; Siegle *et al*. 2014; Cardin *et al*. 2009). Specifically, elevated gamma power in sensory neocortex is related to attentional selection (Fries *et al*. 2001). In the OB, elevated gamma is linked to perceptually demanding discriminations between perceptually similar odors (Beshel *et al*. 2007) and has historically been conceptualized as integral to behavioral states (Martin and Ravel 2014; Eeckman and Freeman 1990). We examined gamma oscillations in the low (40-60 Hz) and high (60-80 Hz) gamma range within each brain structure as rats completed the CAT **(Fig 4**). To do this, we measured the power of gamma oscillations within the hold and odor trial epochs across each task type (odor only, tone attention, switch, and odor attention), and normalized these values to those for odor only trials to highlight the specific contributions of sensory-directed attention as compared to ‘basic’ olfactory discrimination.

**Figure 4.**
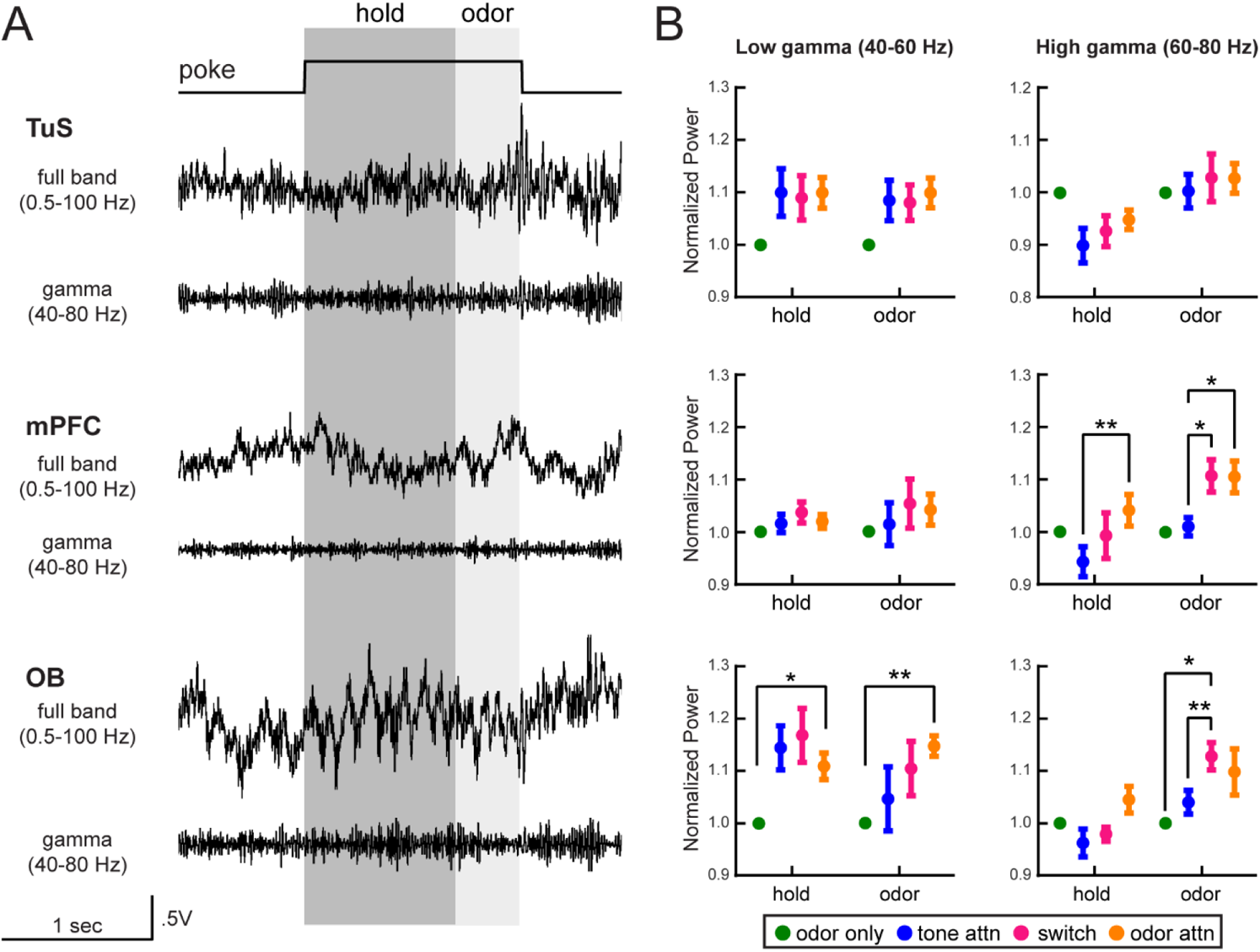
Elevations in gamma power upon intermodal switching and selective attention to odors. **A**. Full band and gamma band filtered (40-80 Hz) traces from the TuS (top) the mPFC (middle) and the OB (bottom) on a single trial of the Carlson Attention Task. Analysis windows for 1 sec hold and 400 ms odor periods are indicated in dark and light gray, respectively. **B**. Quantification of power in the low and high gamma ranges across all task types, normalized to odor only. For each region/frequency band, a 2-way ANOVA with Geisser-Greenhouse correction was completed. TuS, Low gamma: main effect of task type, F(1.62, 6.5)=6.04, p=0.037. mPFC, high gamma: Main effect of trial epoch, F(1.85, 7.38)=13.66, p=0.004. Interaction between trial epoch x task type, F(2.36, 9.44)=5.91, p=0.019. OB, high gamma: Main effect of trial epoch, F(1.38, 5.52)=9.47, p=0.02. Main effect of task type, F(2.22, 8.88)=7.60, p=0.011. n=5 rats, 4.6 +/− 0.5 sessions per rat. On all graphs, asterisks indicate results from Tukey’s multiple comparisons *p<0.05, **p<0.01. All error bars represent SEM.

In the TuS, we observed a slight enhancement in low gamma oscillations with increased attentional demand, but this did not reach statistical significance across rats **(Fig. 4B)**. Interestingly, we observed that high gamma oscillations in the mPFC were elevated during odor attention compared to tone attention **(Fig. 4B)**. We were surprised to observe this enhancement specifically for odor-directed attention in the mPFC, since one might anticipate increased gamma power with increased cognitive demand regardless of sensory modality. In the OB, we observed increased power of low gamma oscillations during odor attention as compared to odor only, indicating that increased attentional demand alone is enough to modify odor information at the earliest stage of processing in the brain **(Fig. 4B)**. Finally, OB oscillations in the high gamma range were elevated in power during the attentional switch relative to odor only and tone attention, suggesting network activity related to cognitive flexibility. While we did uncover some changes in beta band power **(Fig. S2)**, these were not as dramatic across attentional states as was the case with gamma. Overall, these findings indicate changes in local network dynamics in the OB and the mPFC during selective attention to odors, suggesting that attention may modulate odor processing at its most early processing stage (the OB).

### Olfactory bulb gamma oscillations couple with theta phase during selective attention

In some brain regions, the amplitude of high frequency oscillations are structured by the phase of low frequency oscillations, a phenomenon known as phase amplitude coupling (PAC) (Tort *et al*. 2010; Bragin *et al*. 1995; Jensen and Colgin 2007; Lakatos *et al*. 2008; Canolty *et al*. 2006), which is considered a mechanism for attentional selection (Schroeder and Lakatos 2009b). In the OB, PAC between high gamma and respiratory theta becomes stabilized as mice become proficient at discriminating between odors (Losacco *et al*. 2020a), indicating that learning and experience modulates OB PAC. This plus our finding of elevated high gamma in the OB during odor directed attention led us to investigate whether the OB network may engage in PAC during olfactory attention.

To address this, we first computed comodulograms to identify high frequency oscillations coupled to theta phase within the rat OB during the CAT, which revealed strong coupling between theta and high gamma (**Fig 5A**), and much weaker coupling between theta and beta (**Fig S3**). To investigate the significance of theta-high gamma PAC, we examined the trial-by-trial amplitude of high gamma power as a function of theta phase, which indicated high coupling throughout individual sessions and across cognitive states **(Fig. 5B-C)**. Peak phase angle was consistent even comparing correct vs. incorrect trials **(Fig. 5D)**. For each task type, we computed the modulation index (MI) of the PAC, which is a measure of the extent to which a given high-frequency oscillation is structured to a low frequency carrier oscillation. MI values can range from 0.005-0.03 from the hippocampus (Tort *et al*. 2010) and OB (Losacco *et al*. 2020b), and we observed a similar range herein. The distribution in **Fig. 5B** shows the MI theta-high gamma for an example session, and indicates strong PAC. While some individual rats showed modulation of the MI with cognitive state, across the population there were no systematic changes in theta-high gamma PAC with different attentional states **(Fig. 5E)**. Because decreased variance in the peak phase angle in the OB is associated with olfactory learning in a go no-go task (Losacco *et al*. 2020a), we quantified this as well, but did not observe any differences across cognitive states **(Fig. 5E)**. Thus, while the tightly structured theta-high gamma PAC in the OB does not change in magnitude across attentional states, its stability suggests that it possibly supports expert, flexible cognitive function.

**Figure 5.**
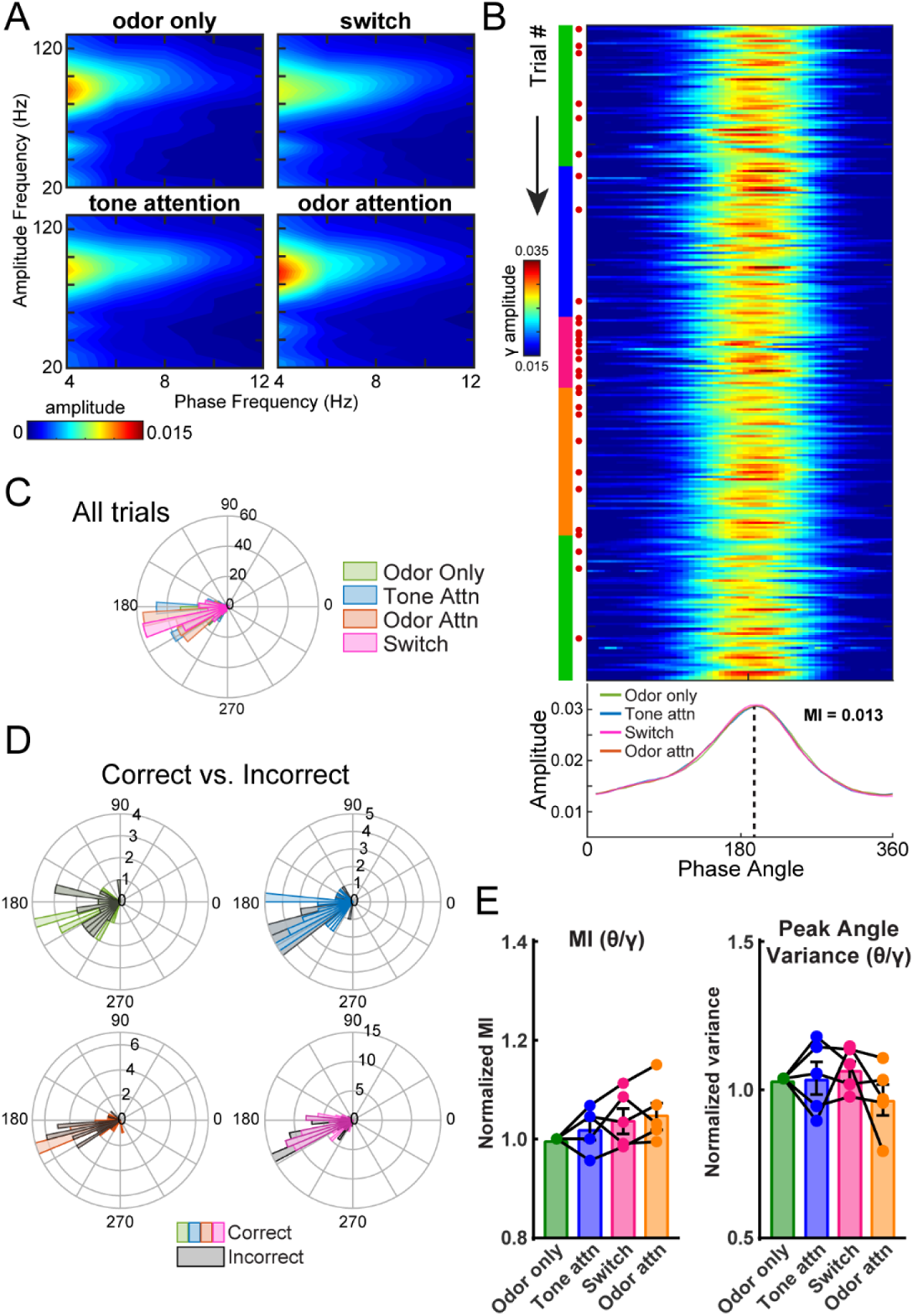
Olfactory bulb gamma oscillations couple with theta phase during selective attention. **A**. Mean comodulograms across rats showing strong coupling between high gamma and respiratory theta frequencies. n=5 rats, 4.6 +/− 0.5 sessions per rat. **B**. Trial by trial theta-gamma for one example session. Green, blue, pink, and orange markings on left side indicate current task type (odor only, tone attention, switch, and odor attention, respectively). Red dots indicate incorrect trials, which expectedly increase in frequency upon switch. The mean amplitude for the session, by task type, is plotted below. MI for the entire session = 0.013. **C**. Polar histogram of peak phase angles by task type for all trials across all sessions for an example rat (n=4 sessions). All task types indicated significant periodicity (Rayleigh test, odor only p<1e118, tone attn p<1e-24, switch p<1e-27, odor attn p<1e-31), and similar distributions (Kolmogorov-Smirnov tests, all comparisons p>0.05). **D**. Polar histograms of correct and incorrect trials for each task type. For this example rat, incorrect trials were pooled across sessions and compared to a randomly-selected equal number of correct trials. Peak phase angle distributions were statistically similar between correct and incorrect trials. (Kolmogorov-Smirnov tests, all comparisons p>0.05). **E**. Left, theta-gamma MI across rats, normalized to MI for odor only trials. One-way ANOVA, F(2.37, 9.48) = 1.96, p=0.019. Right, theta-gamma peak angle variance across rats, normalized to peak angle variance for odor only trials. One-way ANOVA, F(2.03, 8.10)=1.07, p=0.387. n=5 rats, 4.6 +/− 0.5 sessions per rat.Error bars represent SEM.

### Beta oscillations are more coherent between the mPFC and olfactory regions during an intermodal attentional shift

We next tested whether spectral activity between the mPFC and olfactory system might become more coherent during attention. We observed enhanced coherence in the beta range (15-35 Hz) between the mPFC and the TuS specifically during the switch blocks – when the rule has been changed from tone to odor attention, but the rats have not yet successfully switched their attention **(Fig. 6A)**. For the mPFC-TuS, this elevation was specific to the 1 second hold period prior to odor onset **(Fig. 6B)** which corresponds to anticipation. Between the mPFC and the OB, we similarly observed increased coherence in the beta band, but during both the hold and odor epochs **(Fig. 6C-D)**. In contrast, no changes in coherence between the OB and TuS were uncovered (data not shown). Overall, these data indicate that mPFC engagement with olfactory structures is upregulated during a cognitively demanding switch from auditory to olfactory selective attention, suggesting a role for the mPFC in attention-dependent odor processing.

**Figure 6.**
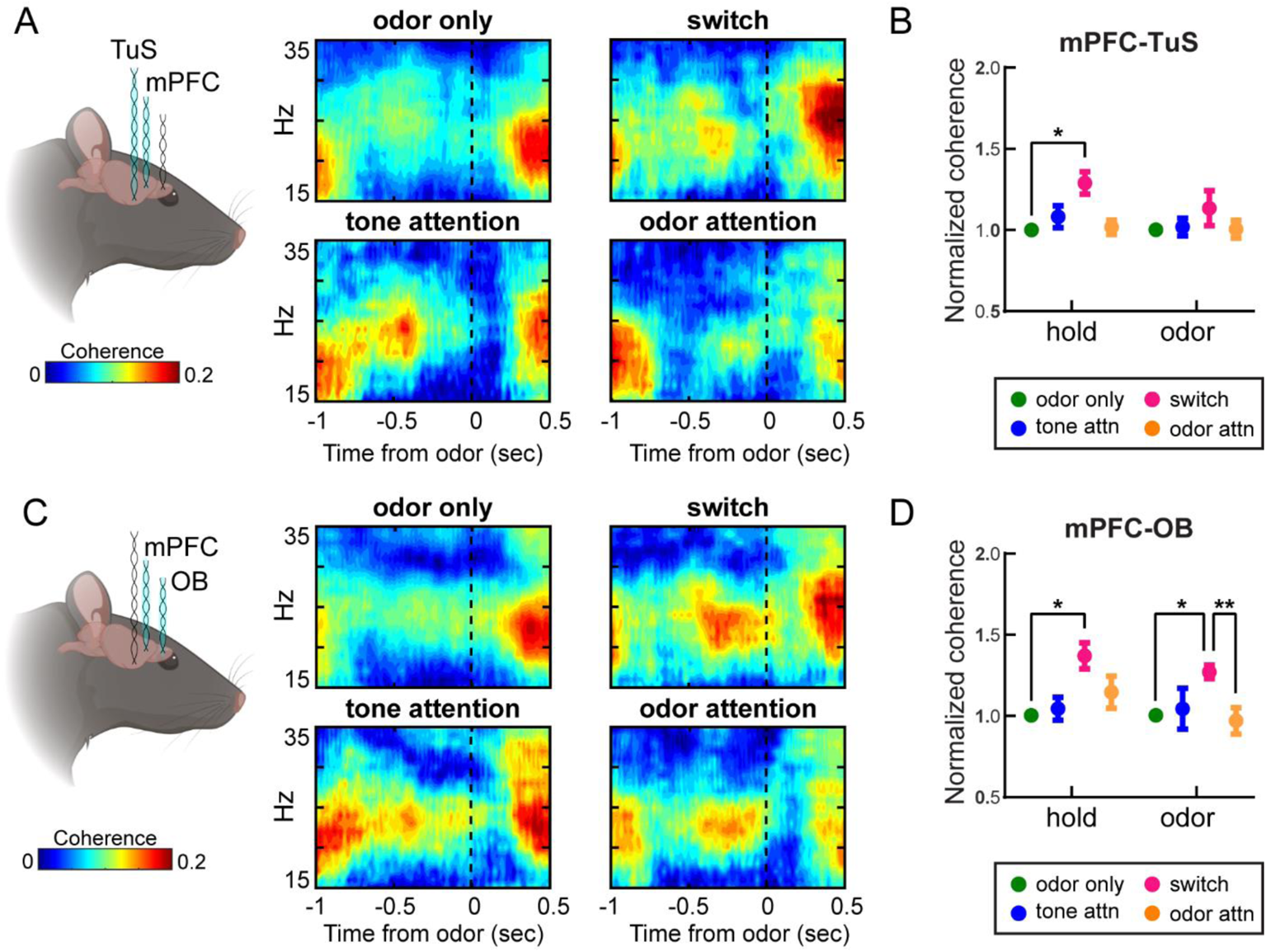
Beta oscillations are more coherent between the mPFC and olfactory regions during intramodal attentional shifts. **A**. Coherogram showing coherence between the mPFC and TuS in the beta range (15-35 Hz) for one example rat across task types (n=3 sessions). Nose poke begins at −1 sec, dotted line indicates odor onset. **B**. Means for OT-mPFC beta coherence across all rats, normalized to odor only. n=5 rats, 4.6 +/− 0.5 sessions per rat. 2-way ANOVA with Geisser-Greenhouse Correction, main effect of task type, F(1.38, 5.41)=6.54, p=0.041. Error bars represent SEM. **C**. Coherogram showing coherence between the mPFC and the OB in the beta range (15-35 Hz) for one example rat across task types. Nose poke begins at −1 sec, dotted line indicates odor onset. **D**. Means for OB-mPFC beta coherence across all rats, normalized to odor only. n=5 rats, 4.6 +/− 0.5 sessions per rat. 2-way ANOVA with Geisser-Greenhouse correction, main effect of task type, F(1.79, 7.15)=13.06, p=0.005. Error bars represent SEM.

### OB-mPFC coherence in the respiratory theta range is strongly upregulated during an intermodal attentional shift to odor attention

Respiration, including fast investigatory sniffing, may structure theta oscillations in not only olfactory regions like the OB (*e.g.,* (Adrian 1942; Kay and Laurent 1999; Buonviso *et al*. 2003)) and TuS (Carlson *et al*. 2014), but also the PFC (Moberly *et al*. 2018; Tort *et al*. 2018b; Biskamp *et al*. 2017; Bagur *et al*. 2021; Zhong *et al*. 2017). Slow wave respiratory theta may serve as a carrier for synchronizing brain regions (Colgin 2013; Fontanini and Bower 2006). This is particularly relevant in an odor-guided task, where correct performance depends upon sampling of the odors via sniffing. We observed that during the attentional switch, there was a striking increase in coherence in the theta range (2-12 Hz) compared to the odor attention state **(Fig. 7A-B)**. While a slight increase was observed during the hold epoch **(Fig. 7A-B)**, this increase was much more pronounced and statistically significant during the odor sampling period **(Fig. 7A-B)**. While our prior analyses had been restricted solely to correct trials for odor only, tone attention, and odor attention, we included correct and incorrect trials for all switch blocks, since this switch state is defined by poor performance and behavioral flexibility, and also because this allowed for the inclusion of comparable numbers of trials in the analysis (see Methods). Thus, we separated trials for switch blocks only into correct and incorrect trials, discarding a random selection of correct trials to match the number of incorrect trials available. This revealed, counterintuitively, a greater coherence on incorrect compared to correct trials **(Fig 7C)**, suggesting that OB-mPFC theta band coherence is upregulated in contexts where behavioral flexibility is required.

**Figure 7.**
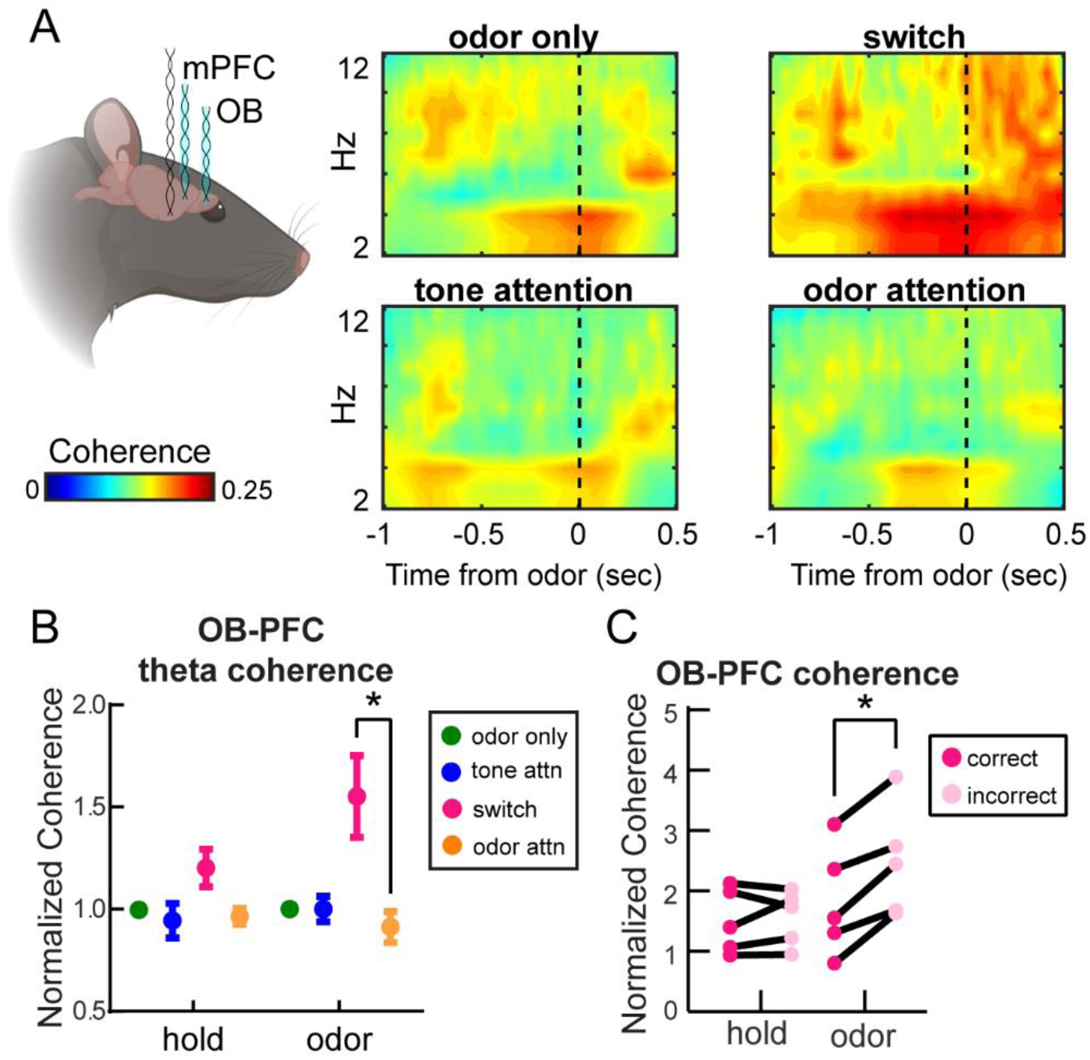
Olfactory bulb and medial prefrontal cortex coherence in the respiratory theta range is strongly upregulated during an intramodal attentional shift to odor attention. **A**. Mean coherogram across rats showing coherence in the theta range (2-12 Hz) across task types. **B**. Mean theta coherence across rats for each trial epoch and each task type. Odor only, tone attention, and odor attention trials include only correct trials from criterion performance blocks. Switch quantification includes all trials from blocks below criterion performance. 2-way ANOVA with Geisser-Greenhouse correction, main effect of task type F(1.3, 5.2)=7.07, p=0.039. Asterisk on graph indicates results from Tukey’s multiple comparison’s test, *p<0.05. Error bars represent SEM. **C**. Theta coherence for correct and incorrect trials during the switch. While there were more correct than incorrect trials, randomly selected correct trials were excluded from this analysis to match the number of incorrect trials. n=5 rats, 4.6 +/− 0.5 sessions per rat. 2-way ANOVA with Geisser-Greenhouse correction, main effect of outcome F(1,4)=76.81, p=0.0009. Interaction between outcome and trial epoch F(1.56, 6.24)=5.39, p=0.048. Asterisk on graph indicates results from Sidak’s multiple comparisons test, *p<0.05.

### Rats maintain highly-stereotyped sniffing strategies despite increased attentional demands

Sniffing behavior in rodents is influenced by many factors including wakefulness, the stimulus being sampled, and motivational state (Clarke and Trowill 1971; Ikemoto and Panksepp 1994; Wesson *et al*. 2008; Kepecs *et al*. 2007; Rojas-Líbano and Kay 2012; Lefèvre *et al*. 2016). As discussed above, whether passive (respiration) or active (sniffing), this behavior subsequently shapes neural activity throughout the brain (Adrian 1942; Macrides *et al*. 1982; Vanderwolf 1992; Colgin 2013; Fontanini and Bower 2006; Verhagen *et al*. 2007; Buonviso *et al*. 2003; Carey *et al*. 2009; Jordan *et al*. 2018; Shusterman *et al*. 2011; Sobel and Tank 1993; Spors *et al*. 2006). We reasoned that if rats adjusted their sniffing when faced with the demand to selectively attend to odors, this could potentially account for the changes in OB-mPFC theta coherence. No prior work has assessed sniffing strategies of rodents during olfactory selective attention. To test this, we trained a separate cohort of rats to perform the CAT before implanting thermocouples in their nasal cavities, allowing us to monitor sniffing behavior by measuring temperature changes (airflow) within the nasal cavity **(Fig. 8A-B)**. Unlike humans, who respond to changing attentional demands by modifying both the depth and timing of respiration (Plailly *et al*. 2008; Arabkheradmand *et al*. 2020), rodents most dramatically employ changes in sniffing frequency during odor active sampling (Cenier *et al*. 2013; Wesson *et al*. 2009; Kepecs *et al*. 2007). Therefore, we quantified sniffing frequency specifically during the hold and odor periods. As illustrated by the example session from one rat in **Fig. 8C**, we observed no clear changes in sniffing behavior across task types **(Fig. 8C-D)**. Instead, the rats displayed a highly stereotyped pattern of sniffing behavior, suggesting that reaching high proficiency on the CAT results in their development of a sensorimotor program that is implemented on each trial, regardless of current attentional demand (**Fig. 8C**, bottom). This was the case across all rats. Although we observed a slight decrease in sniffing frequency specifically during the hold period throughout a session on average (**Fig. 8D, F**), this was confined to the hold period as the rats anticipated odor arrival and sniffing frequency during the odor sampling period remained remarkably constant throughout the sessions for 2/3 rats (**Fig. 8D, F**). Given the lack of changes in sniffing frequency by task type, we investigated whether the timing of sniffs during the odor period were more intentional in relation to odor onset when animals were faced with attending to odor. We examined the time to the first sniff across task types and found no difference, suggesting that sniff timing relative to odor onset is independent of attentional demands **(Fig. 8E)**. We also examined sniffing frequency on correct versus incorrect trials during the switch blocks **(Fig. 8G)** yet did not identify differences in sniffing frequency, providing further evidence that changes in sniffing behavior do not likely account for modulations in OB-mPFC coherence that we uncovered, which were correlated with trial outcome. Together, these data indicate that sniffing strategies in rats are resilient to enhanced attentional demand, providing evidence for covert (rather than overt) olfactory attention in rodents.

**Figure 8.**
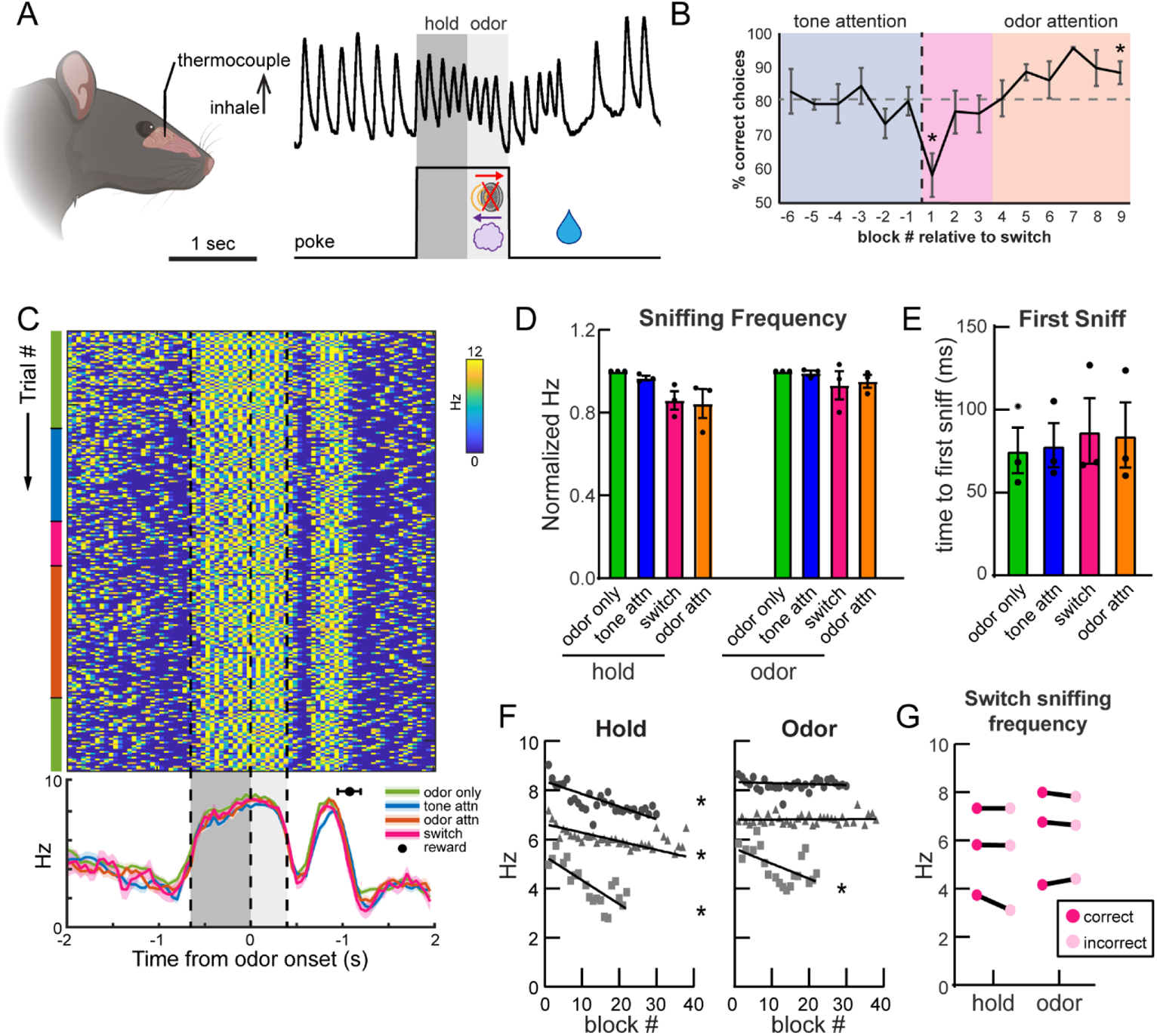
Rats maintain highly stereotyped sniffing strategies despite increased attentional demands. **A**. Sample trace of thermocouple signal from rat nasal cavity on a single trial. 600 ms hold and 400 ms odor epochs are indicated by dark and light gray shading, respectively. **B**. Behavioral performance. Block -1 vs. 1 paired, two-tailed t-test, p = 0.049. Block 1 vs. 9 paired, two-tailed t-test, p=0.026. **C**. Instantaneous sniff frequency for one session, from one example rat (rat 137). Colored bars on the left-hand side indicate the current task type. Dotted lines and light and dark gray shading represent hold and odor epoch respectively. Mean sniffing frequency ± SEM for each task type is plotted below, with the black circle indicating the mean time of reward acquisition (±SEM). **D**. Sniffing frequency means within each trial epoch. 2-way ANOVA with Geisser-Greenhouse correction, main effect of trial epoch F(1,2) = 29.47, p=0.032. No main effect of task type, F(1.04, 2.09)=3.2, p=0.21. Error bars represent SEM. **E**. Time to the first sniff following odor onset. One-way ANOVA with Geisser-Greenhouse correction, F(1.19,2.38)=1.91, p=0.29. Error bars represent SEM. Similar results were seen when calculating time to second and third sniffs, as well as intervals between them (data not shown). **F**. Correlations between block # of the session and sniffing frequency. Rat 137 (circles) hold: R^2^=0.33, F(1,111)=54.46, p<0.0001, odor: R^2^=0.009, F(1,111)=1.03, p=0.31. Rat 138 (squares) hold: R^2^=0.61, F(1,18)=28.61, p<0.0001, odor: R^2^=0.32, F(1,18)=8.406, p=0.009. Rat 139 (triangles) hold: R^2^ =0.3, F(1,168)=73.14, p<0.0001, odor: R^2^=0.0004, F(1,168)=0.07, p=0.79. **G**. Sniffing frequency for correct and incorrect trials in switch blocks only. 2-way ANOVA with Geisser-Greenhouse correction, main effect of trial epoch, F(1,2)=57.65, p=0.017. No main effect of trial outcome. n=3 rats, 3.5 ± 2.5 sessions per rat.

## Discussion

Here we used anatomical, behavioral, and physiological approaches to demonstrate integration of the mPFC with the olfactory system in the context of selective attention to odors. We show that mPFC neurons in the PrL and IL subregions directly and preferentially target the mTuS compared to other olfactory regions, suggesting that they are well-positioned to exert influence on olfactory processing via the mTuS. We then used a physiological and behavioral approach to demonstrate local and interregional effects of attention on network activity within and between the mPFC and olfactory regions, including the OB and TuS. Finally, we found that olfactory sampling behavior is resilient to attentional demand, indicating that olfactory attention may be an “covert” rather than “overt” process. Together, this work adds to a growing body of literature on the possible mechanisms underlying cognitive modulation of olfactory processing and thus perception.

### Insights into mPFC connectivity with the olfactory system

Our tracing experiments uncovered previously unappreciated aspects of mPFC connectivity with the olfactory system. We found that the PrL and IL most densely innervate the mTuS compared with other olfactory regions. By using a combinatorial AAV approach, where Cre expression driven by the CaMKII promotor permits expression of synaptophysin-eGFP/-mRuby, we were able to identify this pathway as excitatory while confidently attributing fluorescence in the TuS (and PCX) to synaptic terminals (primarily in layers 2 and 3) rather than fibers of passage **(Fig. 1)**. We further demonstrated that among PFC subregions, the PrL and IL provide the most projections to the TuS, with the MO coming in third, and these projection neurons mostly reside in layer 5 **(Fig. 2)**. Our data are in agreement with earlier tracing work which established that rat mPFC neurons project throughout the brain, including in the TuS (Vertes 2004), and a recent review proposing that the PrL, IL and MO be grouped together as the ventromedial PFC, based upon their connectivity (Le Merre *et al*. 2021). Our results expand upon previous literature, which used anterograde phaseolus vulgaris-leucoagglutinin tracing (Vertes 2004), by contributing (1) specificity and certainty regarding the specific layers of mPFC→TuS synapses and (2) clarity about at least one mPFC cell type. mPFC glutamatergic neurons modulate their firing during sustained attention (Kim *et al*. 2016) and encode task rules during an intermodal attention task (Rikhye *et al*. 2018), suggesting that TuS-projecting glutamatergic mPFC neurons are positioned to influence olfactory processing during attentionally-relevant behavioral events. Future work illuminating the more specific identities of these TuS projecting mPFC neurons (e.g., via transcriptomics) will be important for disambiguating their specific circuitry and possible contribution to olfactory attention.

### Prefrontal-olfactory network engagement during selective attention

Our multisite LFP recordings during attentional performance uncovered many changes in network activity which expand our appreciation for how the olfactory system is shaped by cognitive state. There are several especially notable outcomes we discuss here.

First, while we know that the mPFC is crucial for attention, no studies have monitored mPFC network activity during olfactory attention, leaving a major void in our understanding of how the mPFC engages with the olfactory system. Because the mPFC is integral for some forms of attention, we predicted it may be recruited during olfactory attention. In support of this, we observed elevated gamma power in the mPFC during odor-directed attention relative to tone attention **(Fig. 4)**. The mPFC is certainly not an olfaction-specific structure, though it is engaged by odor-guided tasks requiring learning (Wang *et al*. 2020) and high working memory capacity (De Falco *et al*. 2019). This elevation in gamma power does not likely reflect increased reward confidence, as behavioral performance was comparable across task types **(Fig. 3C)**. It is interesting to consider whether the mPFC of rodents is predisposed to favor and prioritize olfactory information more so than other sensory stimuli. Nevertheless, these findings exhibit engagement of the mPFC during odor-directed attention, providing support for its inclusion in an olfactory attention network.

### Olfactory attention enhances power of OB gamma oscillations

Our work is the first to monitor OB activity during selective attention. We found that attention powerfully shapes OB activity, which implies that odor information received by structures downstream from the OB, including the TuS, is subject to attention-dependent modulation. Specifically, we observed increased power of low gamma oscillations (40-60 Hz) in the OB during odor-directed attention as compared to odor only discriminations **(Fig. 4)**. Additionally, we observed elevated power of high gamma oscillations (60-80 Hz) while rats attempted to switch their attention from tones to odors **(Fig. 4)**. While increased gamma power in the OB has been associated with successful discrimination of perceptually similar vs. dissimilar odors (Beshel *et al*. 2007), our findings indicate that similar effects can be observed when the odor discrimination is simple/coarse, but the attentional demand is high. Interestingly, elevated low gamma power during odor attention was evident during both the hold and odor epochs, while elevated high gamma power during switch blocks was isolated to the odor sampling period (**Fig. 4B)**. High and low gamma oscillations are considered distinct phenomena in the OB, and are believed to have mechanistically unique origins (Kay 2003), so it is perhaps not surprising to observe modulation of these frequency bands during different attentional demands. Low gamma oscillations are believed to arise from inhibition between local interneurons, are unstructured relative to the sniff cycle, and are functionally mysterious, though they have been observed in states of engaged quiescence (Kay 2003). Our data thus support a potential role for low gamma oscillations in attentionally demanding odor discriminations, though future work is needed to fully appreciate the mechanisms of this.

In contrast, high gamma is structured to the sniff cycle, and is generated by local excitatory-inhibitory interactions (Schoppa 2006; Neville and Haberly 2003; Halabisky and Strowbridge 2003; Lepousez and Lledo 2013). Disruption of high gamma oscillations in the OB impairs odor discrimination, suggesting their importance for basic aspects of odor perception (Lepousez and Lledo 2013). While the mechanisms by which they may be modulated are unclear, one compelling proposition is that neuromodulators, including acetylcholine and noradrenaline, influence excitatory-inhibitory interactions in the OB (Kay *et al*. 2009). This is of particular interest given the role of these neuromodulators in states of attention and arousal (Sara 2009; Yu and Dayan 2005). Our observation that high gamma power elevations during the switch are confined to the odor sampling period is consistent with these mechanistic underpinnings and suggests specific changes in the nature of odor processing as one undergoes a cognitively demanding switch to odor attention.

As mentioned, high frequency gamma in the OB is consistently aligned with the respiratory cycle, which was evident in our PAC analysis **(Fig. 5)**. Recent work demonstrated that OB theta-high gamma PAC is strengthened as mice learn to discriminate odors in a go-no go task, specifically for the go stimulus, suggesting that PAC may support olfactory behavior (Losacco *et al*. 2020a) and leading us to test whether attention employs (or perhaps just simply influences) OB PAC. Our results uncovered highly consistent theta-gamma PAC in the OB across attentional demands, and much weaker coupling between theta and beta oscillations, leading us to focus on theta-gamma PAC. However, we did not observe a decrease in PAC when expert rats completed trials incorrectly (**Fig. 5D)**, suggesting that perhaps PAC is not necessary to successfully discriminate coarse odor pairs, like the ones we used herein. One possible explanation for this difference is that in 2-alternative choice tasks, like the CAT, both stimuli are assigned positive valence, while in go no-go tasks like that used by (Losacco *et al*. 2020a), one stimulus loses positive valence upon learning. Throughout a single session of the CAT, odors temporarily lose their reward-predictive value during tone attention, but it is quickly regained (e.g., **Fig. 3C**). Our data indicate that OB PAC, in rats who have been shaped to expert level on the same odor discrimination over many weeks, is resilient to a temporary lapse in positive odor valence, and perhaps supports flexible behavior supporting attentional switches.

### Beta synchrony integrates mPFC activity within the olfactory network

Our data are the first to show functional coupling between the mPFC and olfactory regions during attentional demands – specifically during an intermodal attentional shift to odors **(Fig. 6)**. Beta oscillations are considered an important mechanism by which long-range communication can occur between brain regions (Spitzer and Haegens 2017; Kopell *et al*. 2000), and further, are implicated in top-down control of attention (Richter *et al*. 2017; Sacchet *et al*. 2015). We observed elevated beta coherence between the mPFC and the TuS as rats attempted to switch their attention from tones to odors (**Fig. 6A-B**). In the context of our finding that the mPFC and the TuS are connected via a unidirectional monosynaptic pathway **(Figs. 1-2)**, these data suggest that communication between the mPFC and TuS is strengthened during attentional shifts. Indeed, ventral striatum-projecting mPFC neurons are implicated in cognitive flexibility by integrating feedback from trial outcomes (Spellman *et al*. 2021), suggesting that the mPFC→TuS pathway could engage in the same processes. Our findings provide further support for the hypothesis that interareal beta oscillations may be a mechanism by which information about behavioral context is conveyed from higher-level cortex to lower-level sensory areas (Bressler and Richter 2015; Wang 2010; Kay and Freeman 1998), and are the first to demonstrate that this concept is applicable to the olfactory system, which possesses unique anatomical organization.

In addition to enhanced mPFC-TuS coherence, we also observed enhanced beta band coherence between the mPFC and the OB **(Fig. 6C-D)**, raising the intriguing possibility that prefrontal influence on olfactory processing could begin as early as the OB. While the OB and mPFC are not connected monosynaptically **(Fig 1)**, they are intermediately connected via bidirectional connectivity with the AON, a pathway known to drive coherence between these structures (Moberly *et al*. 2018). Additionally, these two regions both receive inputs from key neuromodulatory nuclei (Santana and Artigas 2017; Devore and Linster 2012; McLean *et al*. 1989; Rothermel *et al*. 2014; Passetti *et al*. 2000; Devoto *et al*. 2005; Zaborszky *et al*. 1986), which may influence the power of beta oscillations in the OB, perhaps by modifying granule cell excitability (Osinski *et al*. 2018). This raises the possibility that neuromodulators may enable mPFC-OB coherence via simultaneous phasic input to both the mPFC and OB. Cholinergic input increases (Passetti *et al*. 2000; Himmelheber *et al*. 2000) and modulates mPFC firing in the context of attention (Gill *et al*. 2000), and powerfully alters the encoding of odors in the OB (Chaudhury *et al*. 2009; Devore and Linster 2012; Ogg *et al*. 2018; D’Souza and Vijayaraghavan 2014). Indeed, lesioning of cholinergic nuclei results in reduced beta synchrony and increased attentional errors in rats (Ljubojevic *et al*. 2018), indicating at role for cholinergic modulation in attention-related interregional synchrony. Whether cholinergic mechanisms contribute to attention-driven beta synchrony between the mPFC and OB as we observed is an important future question.

### Olfactory sampling is resilient to attentional demand

Odor perception requires the inhalation of an odor, and in rodents this occurs by means of rhythmic inhalation and exhalation of air through the nose in the theta rhythm (Welker 1964; Youngentob *et al*. 1987; Wesson *et al*. 2008; Kepecs *et al*. 2007). The work herein is the first to investigate the influence of olfactory selective attention on sniffing behavior in a rodent. This question is of great interest and importance, since theta oscillations in both olfactory regions and beyond are profoundly shaped by respiration (Adrian 1942; Macrides 1975; Vanderwolf 1992; Tort *et al*. 2018a; Colgin 2013; Zhang *et al*. 2021; Fontanini and Bower 2006; Kay and Laurent 1999; Buonviso *et al*. 2003; Miura *et al*. 2012). Especially relevant to our work, low frequency respiration during freezing behavior can drive strong coherence between OB and mPFC activity, further supporting functional connectivity between these networks (Moberly *et al*. 2018; Bagur *et al*. 2021).

Rodents structure their sniffing in manners influenced by motivation, behavioral task structure, and the sensory stimulus itself (Wesson *et al*. 2008; Kepecs *et al*. 2007; Clarke and Trowill 1971; Ikemoto and Panksepp 1994; Rojas-Líbano and Kay 2012; Lefèvre *et al*. 2016). While some findings suggest that sniffing strategies do not change in the face of increased perceptual difficulty (Wesson *et al*. 2009; Uchida and Mainen 2003), we hypothesized, based on our finding of increased theta synchrony between the OB and mPFC **(Fig. 7),** that enhanced attentional demand may influence sampling strategy. For instance, a rat might increase sniffing frequency during odor sampling when attention to odors versus attending to tones. We found that rats’ sniffing strategies were remarkably resilient to shifting attentional demands, remaining stereotyped as rats flexibly switched their attention from the auditory to olfactory modality **(Fig. 8)**. This is in contrast to some findings in humans, indicating that humans alter the timing and depth of their inhalations during odor anticipation and/or attention (Arabkheradmand *et al*. 2020; Plailly *et al*. 2006). Interestingly, humans also structure their inhalations relative to task structure even when the task is not olfactory in nature, pointing to a role for respiration in structuring and supporting behavioral performance overall (Perl *et al*. 2019). This idea, along with our findings, together raise the intriguing possibility that rhythmic sniffing may even enhance perception of other stimulus modalities (*e.g.,* auditory), perhaps via cross-modal entrainment (Bauer *et al*. 2021; Lakatos *et al*. 2019).

It is interesting to consider sensory sampling via sniffing as analogous to saccadic eye movements (Uchida *et al*. 2006), which contribute to rhythmic attentional sampling in the visual system (Fiebelkorn and Kastner 2019; VanRullen 2016). In the visual system, attention is regarded as overt when it is accompanied by saccadic eye movements to a target and covert when the eyes remain fixated on a central point (Posner *et al*. 1980). The investigation of these different modes of attention and their underlying networks has spanned decades (Posner 2016). While these two processes engage similar brain networks (Rizzolatti *et al*. 1987; Corbetta 1998), suggesting that they may not actually be separate, other work suggests different populations of neurons within these networks may support each type of attention (Thompson *et al*. 2005). Analogously, our observation of covert olfactory attention (*i.e.* olfactory attention that occurs in the absence of attention-specific changes in sniffing behavior; **Fig. 8**) does not necessitate that olfactory attention is *always* covert (*i.e.* sniffing is unaffected), and perhaps different behavioral contexts might engage different olfactory attentional frameworks.

## Conclusion

Taken together, our data support a model of olfactory attention in which the mPFC integrates with olfactory regions at early (OB) and later (TuS) stages of odor processing to form an olfactory attention network. This network encompasses local attention-dependent changes in activity within the OB and mPFC, as well as strengthening of interregional coupling between the mPFC-OB and mPFC-TuS. Our data suggest that changes in sniffing do not drive these effects, highlighting that odor-directed attention, at least in this context, is orchestrated by top-down mechanisms, as opposed to ‘bottom-up’ influences (from odor sampling). Overall, these findings begin to reveal an olfactory attention network and bring us closer to understanding how the brain affords the ability to selectively attend to odors.

## Materials and Methods

### Animals

Adult, male Long-Evans rats were obtained from Charles River (Wilmington, MA) and Envigo (Indianapolis, IN) and maintained in the University of Florida vivarium on a 12:12 light:dark cycle, with food and water provided *ad libitum* until water restriction for behavioral shaping began. All experiments were conducted in accordance with NIH guidelines and were approved by the University of Florida Institutional Animal Care and Use Committee.

### Surgical procedures

For all surgical procedures, rats were maintained on 4-1% isofluorane in 1.5 L/min O_2_ and placed in a stereotaxic frame. The scalp was shaved and cleaned with betadine and 70% ethanol. Analgesia in the form of meloxicam was administered (5 mg/kg s.c.) and the local anesthetic marcaine (5 mg/kg s.c.) was given prior to the cranial incision. A cranial incision was made and the skin was retracted using hemostats.

For viral injections, a craniotomy was then drilled over the region of interest, and a glass micropipette containing AAV was slowly lowered into region of interest. For anterograde mPFC injections **(Fig. 1)**, 100 nL of a 50/50 mixture of Cre-dependent synaptophysin virus (Ef1α-DIO-Synaptophysin-mRuby in IL; Ef1α-FLEX_Synaptophysin-GFP in PrL; both generous gifts from Dr. Marc Fuccillo, Univ of Pennsylvania) (Herman *et al*. 2016) and CaMKII-Cre virus (pENN-AAV9-CaMKII-Cre-SV40; Addgene, Watertown, MA; 105558-AAV9, titer 1×10^13 vg/mL) was injected into the IL, then the PrL, at a rate of 2 nL/sec. For retrograde mPFC injections **(Fig. S1)**, 200 nL total of AAVrg-hSyn-GFP (Addgene 50465-AAVrg; titer 7×10^12 vg/mL) was unilaterally injected at a rate or 2 nL/sec into the mPFC (100 nL in IL, followed by 100 nL in PrL). For TuS injections, 200 nL of AAVrg-hSyn-GFP (Addgene 50465-AAVrg; titer 7×10^12 vg/mL) was injected unilaterally at a rate of 2nL/sec. In all cases, after waiting 5 minutes, the pipette was slowly withdrawn from the brain, the craniotomy was sealed with dental wax, and the incision was sutured.

For electrode implants, the skull was scrubbed with 3% H_2_O_2_ and covered with a thin layer of cyanoacrylate (Vetbond, 3M). Craniotomies were drilled over each brain area of interest, plus 3 craniotomies for 0-80 stainless-steel screws to aid in anchoring the dental cement. After drilling craniotomies over each brain area of interest, the bipolar stainless-steel electrodes (0.005-in outer diameter, Teflon coated to 0.007-in outer diameter) were lowered into the brain and secured with a small amount of dental cement before moving on to the next electrode. Once all wires were placed and secured, an electrical interface board (EIB) (Open Ephys, Cambridge, MA) fitted with a 32 channel connector (Omnetics, Minneapolis, MN) was lowered over the skull, and the electrode wires were secured to the desired channels using gold pins. After the stainless-steel ground wire was secured to a skull screw with conductive silver paint, the whole assembly was secured with dental cement.

For thermocouple implants (Wesson 2013; Uchida and Mainen 2003), following skull preparation as above, a craniotomy was made in the nasal bone (0.9 mm lateral from midline) and a thermocouple wire was lowered 3 mm into the nasal cavity and secured with dental cement. Then, as for the electrode implants, an EIB with a 32-channel Omnetics connector was lowered over the skull, the thermocouple leads secured with gold pins, and a ground wire secured to a skull screw before the whole assembly was secured with dental cement.

Following surgery, rats were returned to their home cages to recover on a heating blanket. The rats received post-operative analgesia for at least 3 days mixed with a palatable gel (5 mg/kg meloxicam in Medigel, ClearH2O, Westbrook, ME). Electrode implanted rats were implanted prior to the beginning of behavioral shaping. Thermocouple implanted rats were shaped prior to surgery and were allowed full water access for at least 24 hours prior to surgery. All rats were allowed to recover for at least 5 days before beginning or restarting water restriction.

### Perfusion and histology

For anterograde mPFC viral injections **(Fig. 1)**, rats were perfused 2-4 weeks following injection. For retrograde mPFC **(Fig. S1)**, and TuS viral injections **(Fig. 2)**, rats were perfused 2 weeks following injection. All rats were overdosed with Fatal-Plus and perfused with cold 0.9% NaCl followed by cold 4% formalin. Brains were dissected and stored in 10% formalin in 30% sucrose prior to sectioning. Alternate 40 um sections were collected with a sliding microtome and stored in Tris-buffered saline with 0.03% sodium azide. For electrode implanted rats, sections were mounted on gelatin subbed slides and stained with 0.1% cresyl violet to confirm electrode locations.

### Image acquisition and quantification

Brain areas of interest were identified using the rat brain atlas (Paxinos and Watson 1997). Images were acquired with a Nikon Eclipse Ti2e fluorescent microscope at 20x magnification using a Nikon 16 MP DS-Qi2 monochrome CMOS camera. For all tracing experiments, successful targeting of the desired subregion was confirmed, and injections with spillover into surrounding regions were excluded. For anterograde mPFC injections (**Fig. 1),** if one of the two injections was on target, we analyzed only that region and disregarded the other. Overall, we analyzed 8 rats with on target PrL injections and 5 rats with on target IL injections, with 3 rats having both PrL and IL quantified. From these rats, images for quantification were acquired as follows: for the TuS, 11 images per rat, evenly spanning 2.7mm anterior – 0.8 mm posterior Bregma. For the PCX, 17 images per rat were quantified, evenly spanning 3.7mm anterior – 4.8mm posterior Bregma. For the AON, 6 images per rat were quantified, evenly spanning 5.7mm-2.7mm anterior Bregma. For retrograde TuS injections (**Fig. 2**), 3-10 (6.46±2.48) PrL/IL-containing sections and 1-6 (3.9±1.46) OFC-containing sections were imaged (n = 6 rats).

After acquiring images, ROIs were drawn around each area of interest and fluorescent puncta or cell bodies were detected using semi-automated counting algorithms created within NIS elements software (Nikon) based on their fluorescence intensity and size. Cell or puncta counts were then normalized to the ROI area for comparison across regions. For layer-specific quantification (**Fig. 2E),** custom MATLAB code was used to determine the layer in which each counted cell resided. The medial and lateral TuS were defined as the medial and lateral third of the TuS, to ensure clear separation between the regions. In puncta quantification, we initially differentiated between the anterior and posterior PCX, which was divided based on the presence or absence respectively, of the lateral olfactory tract. Because we observed no differences in puncta between the anterior and posterior PCX for PrL (paired, two-tailed t-test, p=0.32) or IL (paired, two-tailed t-test, p=0.12), we combined them for the data and analyses shown in **Fig. 1**.

### Olfactory and auditory stimuli

The odors used for all experiments were isopentyl acetate and limonene(-), obtained at their highest available purity (Sigma, St. Louis, MO), and diluted in mineral oil (Sigma) to 0.5 Torr so that they possessed equal vapor pressures. Odors were delivered through independent lines via an air-dilution olfactometer at 2 L/min via a custom 3-D printed nose-poke port. The auditory stimulus was a 2.5 kHz tone (∼70dB) generated with a piezo speaker (RadioShack, Boston, MA).

### Carlson Attention Task

Rats were water restricted to no less than 15% of their initial body weight and were shaped on the Carlson Attention Task (CAT) as described in detail previously (Carlson *et al*. 2018). Briefly, rats were first shaped on single-modality 2-alternative choice (2-AC) tasks in blocks of 20 trials, starting with tone-on/tone-off 2-AC, then odor A/odor B, before learning the multi-modal attention task. In the final task, rats initiated a trial by nose poking in a center port. They were required to hold for 1 second (for LFP rats) or 600 ms (for sniffing rats) before stimulus delivery, and were then required to remain for at least 400 ms for stimulus delivery (the prolonged hold period for the rats contributing LFP data was implemented to provide a sufficient window for subsequent analyses). After leaving the center port, the rats had 4 seconds to make a choice at either the left or right port. Correct choices were rewarded with 15 uL of 2 mM saccharin in water, and incorrect choices were unrewarded. If no choices were made in the 4 second window, the trial was recorded as an omission. After the 4 second window, an additional 1 sec inter-trial interval (ITI) was implemented which was reset by a nose poke during that second. Thus, across all trials the rats were out of the center port for one full second prior to their trial-initiating poke. Sessions began with 3 blocks of odor only, in which there were no competing tone cues. After completing 3 blocks at ≥80% correct, the rats began receiving simultaneous olfactory and auditory cues, and were required to complete 6 blocks of tone attention (attending tones, and ignoring odors) at ≥80% correct. After this, an uncued rule change occurred, requiring the rats to now attend odors, and ignore tones. After 6 blocks at ≥80% correct on odor attention, the rats completed 3 more blocks of odor only at the end of the session. Odor only blocks were included at the beginning and end of the session to neutralize any potential effects of motivation. For all task types, trial combinations were pseudorandomly presented, such that equal numbers of each trial type were given in each block of 20 trials.

For LFP recordings, this shaping process occurred over the course of 33-46 (40.8 ± 2.3) sessions, resulting in expert rats who had switched their attention from tones to odors 8-11 (9.8 ± 0.6) times prior to the final recorded sessions included in our data analysis. Each rat contributed 3-6 (4.6 ± 0.5) sessions of expert performance to the analysis. For sniffing recordings, shaping occurred over 63-72 (67.3 ± 2.4) sessions, rats switched their attention 11-17 (13.6 ± 1.8) times before recorded sessions, and contributed 1-6 (3.5 ± 2.5) sessions of expert performance to the analysis. Shaping with the rats used for sniffing took more sessions for them to reach expert performance since they were delayed in learning/performance due to surgical implantation of thermocouples mid task acquisition.

### Data acquisition

LFPs from all electrodes were digitized using an RHD 2132 headstage (Intan Technologies, Los Angeles, CA), amplified using a PZ5 amplifier (Tucker-Davis Technologies, Alachua, FL), and acquired at 3 kHz using OpenEx and an RZ2 BioAmp processor (Tucker-Davis Technologies). Tethering of the rats to the PZ5 occurred via a flexible ultralight tether with a commutator in-line to allow free movement. Entrances to the left, right, and center ports were detected by infrared beam breaks and acquired at 380 Hz. Behavioral and stimulus delivery events were simultaneously recorded in OpenEx using the RZ2 BioAmp processor, allowing for synchrony between the behavioral and neural events. The thermocouple signals were acquired similarly along with behavior, but using Synapse software with a sampling rate of 610 Hz.

### Local field potential analysis

To minimize potential multisensory influences (Gnaedinger *et al*. 2019), all trials analyzed were tone-off trials **(Fig. 3B)**(Carlson *et al*. 2018), and came from blocks where behavioral performance was ≥ 80% correct. For odor only, tone attention, and odor attention, only correct trials were included in all analysis (unless otherwise specified; **Fig. 5D**). For switch blocks, where the rat is by-definition performing poorly and responding to negative reward feedback, we included correct and incorrect trials. Because only tone-off trials were analyzed, there were twice as many odor only trials compared to tone attention and odor attention. Therefore, we randomly discarded half of the odor only trials, preserving the proportion that were from the beginning/end of the session as well as the proportion of trial types. The data were imported into MATLAB and traces spanning −10 to 8.4 sec from odor onset from each trial were downsampled to 1 kHz, filtered 0.5-100 Hz using a 2^nd^ order bandpass filter, and 59-61 Hz using a 2^nd^ order band-stop filter. These large segments of data were further filtered to avoid edge artifacts from filtering, but smaller segments were used for later analysis. Power and coherence were computed using the Chronux toolbox (Mitra and Bokil 2009)(http://chronux.org), and raw values were normalized to odor only trials, to identify effects specifically related to attentional demand. Specifically, multi-taper power spectra and coherence were computed using 5 tapers, and mean power was determined by averaging within the given frequency range. For power and PAC analyses, a single LFP trace from the bipolar electrode was used. For coherence analyses, a subtracted trade was first created from the bipolar electrode. For power and coherence analyses, theta was defined as 2-12 Hz, beta as 15-35 Hz, low gamma as 40-60 Hz, and high gamma as 60-80 Hz.

PAC analysis was completed using MATLAB routines from Tort et al., 2010, wherein the Hilbert transform method was used to determine the phase of the carrier oscillation (theta), and separately, the envelope of the high frequency oscillation (beta or high gamma). The amplitude of the beta/high gamma fast oscillation across 51 bins of the theta phase was plotted to demonstrate PAC strength **(Fig. 5B)**. Theta was defined as 2-10 Hz, beta defined as 15-35 Hz, and high gamma defined as 65-100 Hz. For each trial, a single OB LFP trace spanning from −3 to +1.5 sec from odor onset was used. A larger segment of time was used for these analyses because more samples were required to examine coupling with theta frequencies down to 2 Hz. Still, this time segment allows for the analysis to be contained to an individual trial without any overlap with neighboring trials.

### Sniffing analysis

As with the LFP analyses, all analyses on the sniffing data were restricted to trials where the tone was off. Additionally, only correct trials were examined for odor only, tone attention, and odor attention blocks, while correct and incorrect trials were examined for switch blocks. The data were imported to MATLAB and traces spanning −10 to 8.4 sec from odor onset from each trial were filtered using a 2^nd^ order band-pass filter from 0.5-10 Hz. After extracting trials as described above for the LFP analyses, these filtered traces were convolved with an 8 Hz Morlet wavelet and peaks detected. Detected peaks were visually inspected and false positives were manually rejected using a custom MATLAB GUI. The resultant peaks were used to calculate the instantaneous sniffing frequency throughout the trial. Then, instantaneous frequencies were averaged within each trial epoch (i.e. hold, odor), and normalized to odor only.

### General statistical methods

Semi-automated routines were used to ensure rigorous data extraction and analyses. Details regarding specific statistical tests can be found in their respective results sections and/or figure legends. Unless otherwise stated, all values are mean ± SEM.

## Acknowledgements

This work was supported by National Institutes of Health grants R01DC016519 and R01DC014443 to DW, R01DA049545, R01DA049449, and R01 NS117061 to MM and DW, R01DC006213 to MM, and F32DC018232 to HC. We thank Dr. Mark Fuccillo for generously sharing reagents.

## Conflicts of Interest

The authors have no perceived or real conflicts of interest to declare.

---

**Supplemental Figure S1.**
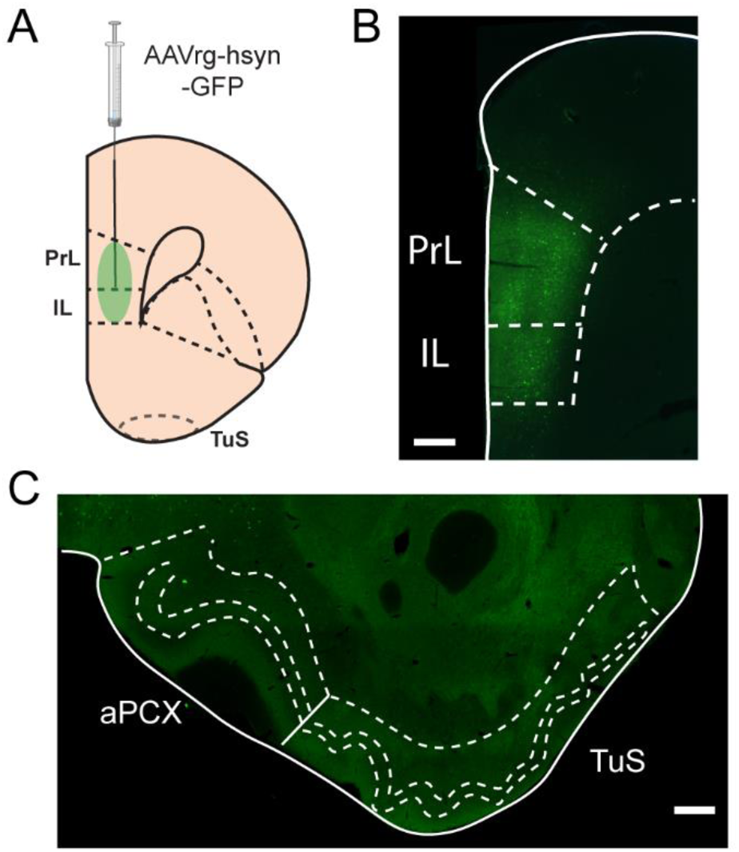
PCX and TuS projection neurons do not innervate the mPFC. **A**. The PrL and IL were injected with AAVrg-hsyn-GFP to identify possible mPFC-projecting neurons. **B**. Example injection site showing spread through the PrL and IL cortex. Scale bar 250 µm. **C**. Example image showing lack of labeled cells in the TuS and aPCX, indicating that these structures do not project to the mPFC. Similar results were seen in 2 rats. Scale bar 250 µm.

**Supplemental Figure S2.**
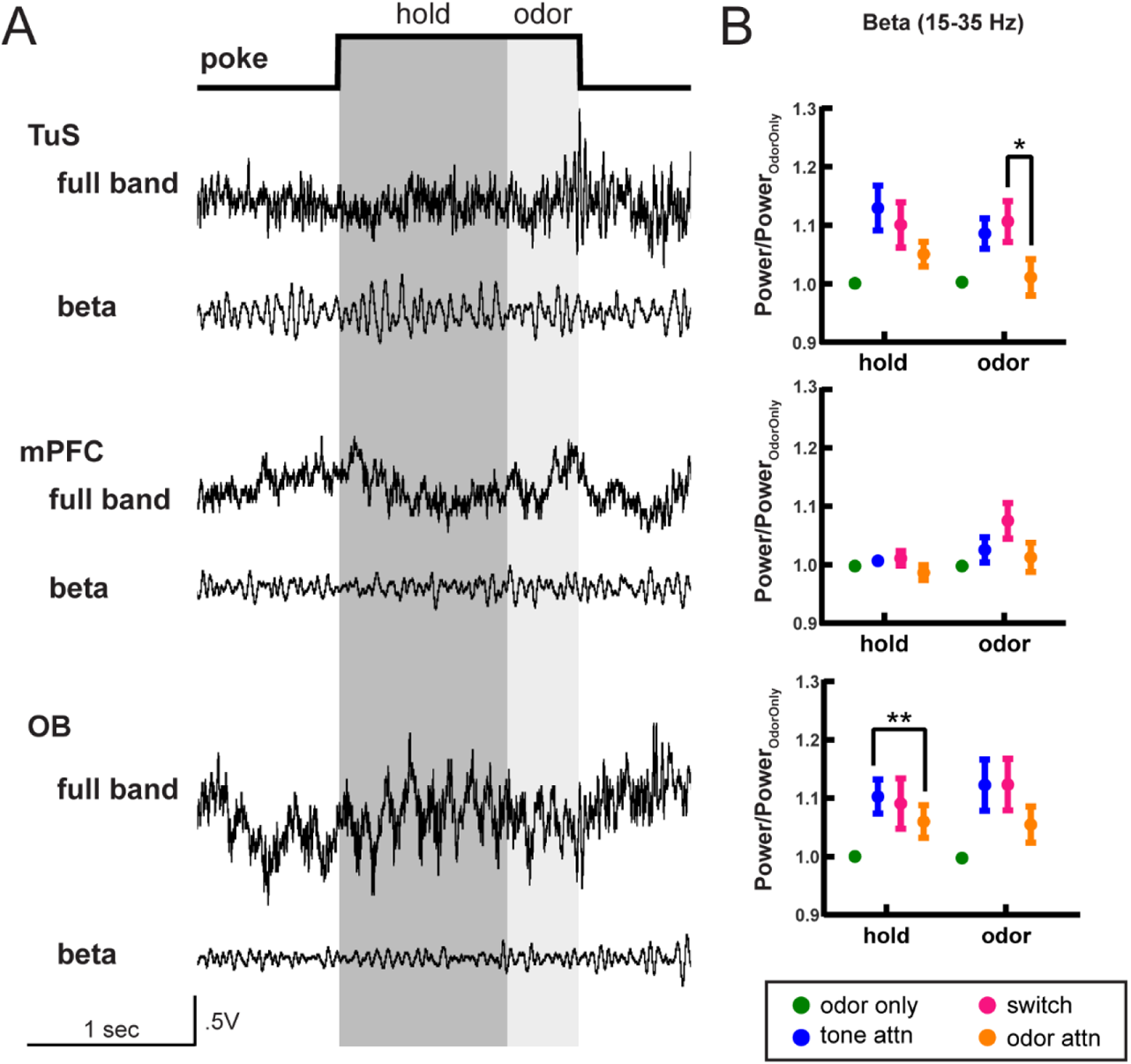
Beta oscillation power during attentional states. **A**. Full band and beta band filtered (15-35 Hz) traces from the TuS (top), mPFC (middle) and OB (bottom) on a single trial of the Carlson Attention task. Analysis windows for hold and odor periods are indicated in dark and light gray, respectively. **B**. Quantification of power in the beta range across all task types, normalized to odor only. Statistical tests were 2-way ANOVA with Geisser-Greenhouse correction. TuS: main effect of task type, F(1.94,7.77)=16.37, p=0.0017. mPFC: main effect of task type, F(2.00,8.02)=4.923, p=0.0402. OB: main effect of task type, F(1.49,5.97)=16.52, p=0.0047. On all graphs, asterisks indicate results from Tukey’s multiple comparisons, *p<0.05, **p<0.01. All error bars represent SEM. n=5 rats, 4.6 +/− 0.5 sessions per rat.

**Supplemental Figure S3.**
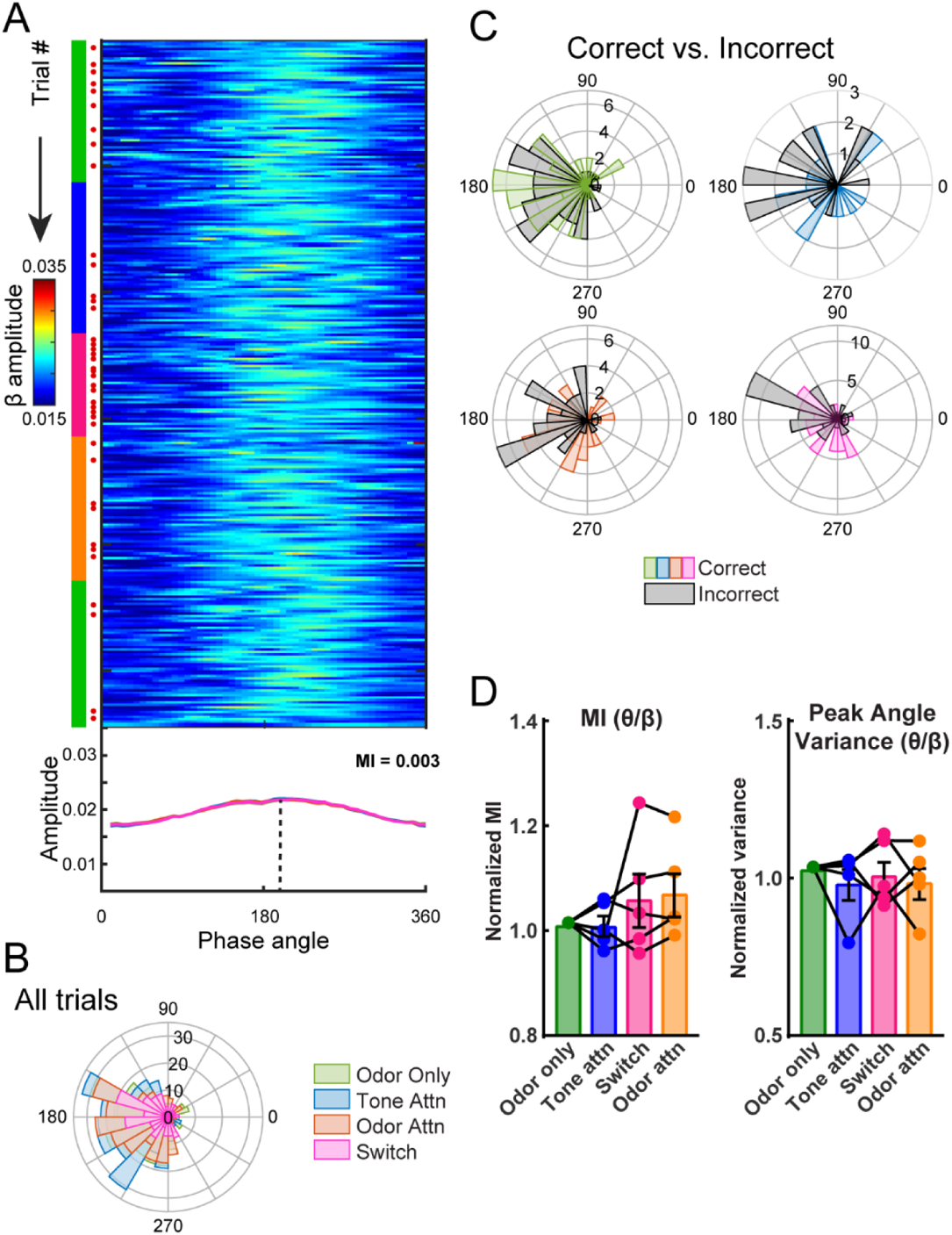
Modest theta-beta phase amplitude coupling in the CAT. **A**. Trial by trial theta-beta PAC for one example session. Green, blue, pink, and orange markings on the left side indicate current task type (odor only, tone attention, switch, and odor attention, respectively). Red dots indicate incorrect trials. The mean amplitude for the session, by task type, is plotted below. MI for the entire session = 0.003. **B**. Polar histogram of peak phase angles by task type for all sessions for this example rat (n=5 sessions). All trial types showed significant periodicity (Rayleigh test, odor only p<1e-27, tone attn p<1e-8, switch p<0.05, odor attn p<1e-12) and similar distributions (Kolmogorov-Smirnov tests, p>0.05 for all comparisons except odor only vs. switch, p=0.046). **C**. Polar histograms showing correct and incorrect trials for each type. For this example rat, incorrect trials were pooled across sessions and compared to a randomly selected equal number of correct trials (n=5 sessions). Peak phase angle distributions were statistically similar between correct and incorrect trials for odor only, odor attention, and switch trials (Kolmogorov-Smirnov tests, p>0.05) but statistically different for tone attention (p=0.041). **D**. Left, theta-beta MI across rats, normalized to MI for odor only trials (One-way ANOVA, F(1.19,4.78)=1.22, p=0.34). Right, theta-beta peak angle variance across rats, normalized to peak angle variance for odor only trials (One-way ANOVA, F(2.18,8.31)=0.35, p=0.72). n=5 rats, 4.6 +/− 0.5 sessions per rat.

## Notes

### Competing Interest Statement

The authors have declared no competing interest.

